# Single-cell sequencing of individual retinal organoids reveals determinants of cell fate heterogeneity

**DOI:** 10.1101/2023.05.31.543087

**Authors:** Amy Tresenrider, Akshayalakshmi Sridhar, Kiara C. Eldred, Sophia Cuschieri, Dawn Hoffer, Cole Trapnell, Thomas A. Reh

## Abstract

With a critical need for more complete *in vitro* models of human development and disease, organoids hold immense potential. Their complex cellular composition makes single-cell sequencing of great utility; however, the limitation of current technologies to a handful of treatment conditions restricts their use in screens or studies of organoid heterogeneity. Here, we apply sci-Plex, a single-cell combinatorial indexing (sci)-based RNA-seq multiplexing method to retinal organoids. We demonstrate that sci-Plex and 10x methods produce highly concordant cell class compositions and then expand sci-Plex to analyze the cell class composition of 410 organoids upon modulation of critical developmental pathways. Leveraging individual organoid data, we develop a method to measure organoid heterogeneity, and we identify that activation of Wnt signaling early in retinal organoid cultures increases retinal cell classes up to six weeks later. Our data show sci-Plex’s potential to dramatically scale-up the analysis of treatment conditions on relevant human models.

## Introduction

Over the past decade, rapid progress has been made towards designing *in vitro* systems that recapitulate the structure and function of organs. These organs in a dish or “organoids’’ now exist for the retina, brain, kidney, intestine and more^1–9^. The extent to which retinal organoids recapitulate development of the human fetal retina is nothing short of remarkable. Retinal organoids not only contain all of the major neuronal and glial (Müller glia) cell classes of the retina, but also mimic the developmental timing and laminar organization of the retina^10^. Importantly, retinal organoids represent an accessible and renewable *in vitro* system for studies of development^11^ and disease progression. When generated from patient-IPSCs, retinal organoids can model patient-specific disease phenotypes^12, 13^, be used for the development of personalized medicine, or possibly provide a source of cells for replacement in degenerative diseases^14–16^.

However, several roadblocks exist that limit retinal organoids from reaching their full potential. These include organoid-to-organoid heterogeneity, batch effects, high costs due to specialized culture reagents with long culture times, and lack of relevant support cells such as immune cells and circulatory cells^17^. While several well-established protocols exist for retinal organoid generation, a common theme across the vastly different differentiation schemes is the requirement for manual selection or dissection of organoids that visually appear retinal^18, 19^. A protocol has been developed which eliminates the need for dissection, but it relies on specialized micro-well plates that have not been widely adopted and it still only produces ∼80% retinal organods^20^. The heterogeneity in these cultures reflects our incomplete knowledge and control of the developmental program that generates retinal cell classes. Sources of such heterogeneity are difficult to define with traditional bulk sequencing methods^21^. Single-cell sequencing analysis allows for the assessment of the milieu of cell classes within a heterogeneous population, but without the ability to associate cells with individual organoids, it’s hard to know how experimental perturbations impact heterogeneity across organoids. Additionally, high levels of variability can mask the impact of perturbations compared to controls. For these reasons, the presence of non-retinal brain cells in retinal organoid cultures, in an unpredictable and heterogeneous manner, limits their reproducibility, interpretability, and utility especially for large-scale experiments^18, 19, 22^.

To advance the standardization and utility of retinal organoids, we develop a high throughput organoid screening technology aimed at identifying culture conditions that produce organoids of desired compositions with greater homogeneity. The most common current techniques for screening in organoids rely on microscopy-based approaches that implement measurements of size and morphology, use viability stains, and/or perform immunostaining/fluorescence for a small number of predetermined marker genes^23–26^. With single-cell sequencing, the ability to detect all expressed genes enables observation of the full array of cell classes in a culture, permits the discovery of unpredictable cellular states, and allows for a treatment’s effect to be measured across all cell classes, not just those traceable with a marker. However, the high per sample cost of the most popular single-cell sequencing platforms can’t practically scale past a handful of conditions (Table S1). sci-Plex, which builds upon single cell combinatorial indexing (sci-RNA-seq), facilitates the labeling and simultaneous processing of thousands of conditions with a single-cell resolution output^27^. Originally performed in cell culture, sci-Plex was recently applied to thousands of individual zebrafish embryos exposed to genetic or temperature-based perturbations^28, 29^. Leveraging the statistical power of individual-embryo single-cell data to detect significant changes in cell class abundance aided in novel developmental findings. Individual organoids have been sequenced at single-cell resolution in a handful of cases, but, despite increasing interest, the experimental designs and level of replication are sharply constrained by the cost and throughput of the droplet-based single cell sequencing technologies^30, 31^. Adapting sci-Plex for individual organoids would increase the capacity to process them at single-cell resolution and promote the development of culturing protocols that faithfully and homogeneously recapitulate *in vivo* cell class compositions (Table S1).

Here we demonstrate the first application of sci-Plex in organoids. To confirm the validity of the method, we show strikingly similar cell class compositions of matched bulk retinal organoids prepared by sci-Plex compared to commercially available droplet-based single-cell sequencing methods after culturing for 28, 78, or 185 days. We then perform sci-Plex on 410 individual organoids while modulating key developmental signaling pathways. We find that the staggered activation of BMP followed by Wnt signaling produces organoids with a greater proportion of retinal cells compared to BMP activation alone without increasing organoid-to-organoid heterogeneity. With the depth of information obtained across hundreds of individuals, we vastly expand the types of analyses possible in single-cell sequencing-based methods in organoids, and we identify a culturing protocol that improves the composition of retinal organoids.

## Results

### Successful multiplexing of retinal organoids using sci-Plex

To demonstrate that multiplexed samples from sci-Plex have 1) cell class compositions comparable to those recovered using commercial droplet-based techniques, and 2) cell class compositions that would be expected developmentally, retinal organoids cultured for 29 (10 pooled), 78 (5 pooled), or 185 (5 pooled) days were dissociated into single-cell suspensions. A subset of the cells were processed using the 10x Chromium platform while the remaining cells were subjected to sci-Plex (**Figure 1A**). For sci-Plex, the cells in each of the three single-cell suspensions were lysed to expose their nuclei, single-stranded and polyadenylated DNA oligos (hashes) with sample-specific barcodes were added to the suspensions, and the hashes were chemically crosslinked to the nuclei by paraformaldehyde (PFA) fixation. All samples were then combined and subjected to 2-level sci-RNA-seq whereby the hashes are further barcoded as if they were mRNA^32, 33^(**Figure S1A**). After processing and filtering, 12,867 cells were recovered by sci-Plex (median UMI for D29: 5512, D78: 5797, D185: 5695.5) and 16,914 were recovered by 10x (median UMI for D29: 6335, D78: 4118, D185: 3950) (**Figures S1B-H**). Using the hash barcodes, the sample from which a cell originated was determined for 95% of the cells (**Figures S1D-E**). The 10x and sci-Plex datasets were then integrated using Seurat’s MNN-CCA methodology^34^, and cell classes were annotated from UMAPs using marker gene expression^35–46^**(Figures 1B-C, S1I-L, Table S2).**

**Figure 1.**
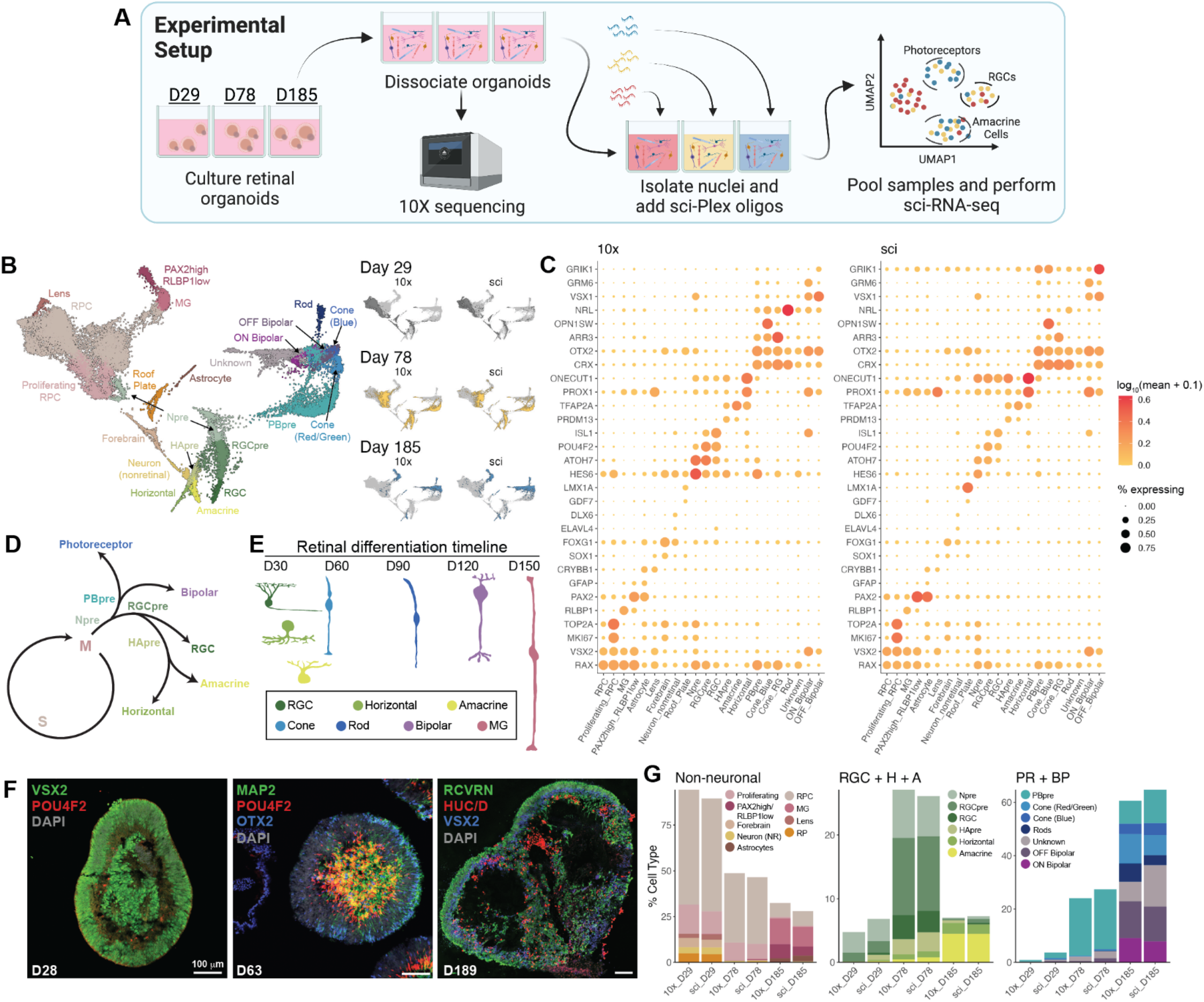
Comparison between sci-Plex and 10X. A) Retinal organoids were cultured for 29, 78, or 185 days before being dissociated into single-cell suspensions. The cells were split and either processed by a standard 10x RNA-seq pipeline or prepared by sci-Plex. In sci-Plex, the cells are lysed in age-specific wells, leaving the nuceli intact. Single-stranded polyadenylated DNAoligos, called “hashes”, are added to each well. The hashes contain well-specific barcode sequences and have an innate affinity for the nuclei which helps recruit the oligos to the surface of the nuclei where we permanently crosslink the hashes to the nuclei. After crosslinking, the nuclei from all wells are pooled and subjected to sci-RNA-seq. B) Integrated UMAP of sci-Plex and 10x datasets derived by Seurat’s CCA-MNN methodology. Left: Cells are colored by cell type. RPC: retinal progenitor cell, MG: Müller glia, Npre: retinal neuronal precursor, RGC: retinal ganglion cell, RGCpre: RGC precursor, HApre: horizontal/amacrine precursor, PBpre: photoreceptor/bipolar precur­sor. Right: Cells are faceted by organoid age and technology. Cells are colored by age. **C)** Dot plot of the genes used to define cell types in B. The size of the dot indicates the percent of cells that express the gene of interest. The color indicates the Iog10 mean UMIs per cell. **D)** A representation of the established view of retinal neuron developmental trajectories. **E)** The timeline of retinal cell differentiation in human organoids. F) Immunostaining of D28, D63, and D189 organoids for progenitor (VSX2-D28), RGC (POU4F2), photoreceptor/bipolar (OTX2), neuronal (MAP2), amacrine/horizontal/RGC (HuC/D), bipolar cells (VSX2-D189) and photoreceptor (RCVRN) markers. Scale bar represents 100 µm. **G)** Using the cell type assignments from the integrated dataset, cell types were split into Brain/Non-neuronal (Non-neuronal), RGC+Horizontal+Amacrine (RGC+H+A), and Photoreceptor+Bipolar (PR+BP) categories. Cells were counted and the % Cell Type was determined for each technology and organoid age.

The retina develops in a highly coordinated way with all retinal neurons and the Müller glia (MG) arising from a shared retinal progenitor cell (RPC) population^47^(**Figure 1D**). Across vertebrates from zebrafish to chick, mice, and humans, the main cell classes and the order in which they develop are conserved. Furthermore, the order and developmental timing is roughly conserved between human retinal organoids and the human retina in the developing fetus^10^(**Figure 1E**). Staining of retinal organoids supported previous observations indicating that the layered organization of the retina is maintained in organoids^10^(**Figure 1F**). We used the knowledge of developmental timing to determine how well sci-Plex segregated the samples of differing ages (**Figures 1B,G** and **S1K**). As expected, Day 29 (D29) organoids included mainly RPCs (VSX2, RAX), differentiating retinal ganglion cells (RGCs: POU4F2, ISL1), and other retinal neuron precursors as well as brain progenitor cells with forebrain (FOXG1, SOX1), neuronal (FOXG1, ELAVL4, DLX6), and roof plate (GDF7, LMX1A, RSPO1) characteristics (**Figure 1C,G** and **S1J-K**). Day 78 (D78) organoids have far more RGCs, horizontal (TFAP2A, PROX1), amacrine (TFAP2A+, PROX1-), and photoreceptor precursor cells (OTX2, CRX) (**Figure 1C,G** and **S1K**). By Day 185 (D185), rods (NR2E3), cones (blue: OPN1SW+, ARR3+, red/green: ARR3+, OPN1SW-), and bipolar cells (OFF: GRIK1, ON: GRM6) were dominant (**Figure 1C,G** and **S1J-K**). Muller glia (MG: RLBP1) and astrocyte (PAX2, GFAP) clusters were also defined (**Figures 1B-C,G** and **S1J-K**). Comparing directly between 10x and sci-Plex, the cell class distribution was highly concordant for most cell classes although some low abundance cell classes were detected more frequently in sci-Plex, possibly due to differences between nuclear and whole cell transcript recovery (**Figure 1G**). We also found that processing just the sci-Plex data using Monocle3 resulted in a comparable depth of cell class classification as the integrated data (**Figure S1L-N**). These analyses demonstrated that sci-Plex can both identify a cell’s well of origin and correctly assign its cell type, underlining the utility of sci-Plex for multiplexing experiments with organoids.

### Detection of cell class abundance changes across treatments through multiplexing of individual organoids

We next aimed to use the multiplexing ability of sci-Plex to screen conditions that modulate signaling pathways in order to 1) better understand the effect signaling pathways have on cell fate decisions in young retinal organoids and 2) use this information to design better organoid differentiation protocols. The BMP, Sonic hedgehog, Wnt, and FGF signaling pathways are all key regulators of eye and brain development^48, 49^. Modulators of these pathways: BMP4 (BMP protein), SAG (Sonic hedgehog agonist), CHIR99021 (Wnt agonist) and SU5402 (FGF inhibitor) have all been used in published retinal organoid protocols^19^. However, there is no agreed upon “best” combination of these factors.

In order to perform a systematic and quantitative statistical analysis, sci-Plex was used to label 314 individual organoids from seven different conditions in which signaling modulators were added at Days 6-12 (BMP4) and/or Days 14-18 (SAG, CHIR99021, SU5402). Organoids were prepared for sci-Plex 28 days after initiation of differentiation (**Figure 2A-B**). At this point in the retinal differentiation protocol, it is standard practice to select for organoids that morphologically appear retinal under brightfield microscopy and discard visibly non-retinal tissues. In our experiments, no selection was applied as we aimed to understand what was driving the heterogeneity that obligates such selection steps. It is because of our desire to better understand the effects of signaling modulation on the choice between a retinal and non-retinal cell fate, that such an early time point was chosen. At this stage, we would expect most of the cells to be retinal progenitors. Additionally, because we were not removing the non-retinal cells from the organoids, we wanted to capture early timepoints in case the retinal cells were negatively affected by the continued presence of non-retinal cells. We distributed the organoids into 96-well plates so each well harbored a single organoid, permitting the hash-based barcoding to be performed on individual organoids. After hashing, cells were subjected to sci-RNA-seq3, a version of combinatorial indexing that greatly increases the throughput of the method^33^. After filtering, 203,511 cells (median UMI: 350) were recovered (**Figures S2A-F**). The median number of cells recovered per individual was 505, however recovery varied by treatment with CHIR treatment (BMP and Wnt activation) returning the highest number of cells per organoid at 870 (**Figure 2C**). Our successful recovery of cells from hundreds of individual organoids set us up to perform statistical analyses relating to cell class abundance and organoid to organoid heterogeneity in an unprecedented manner.

**Figure 2.**
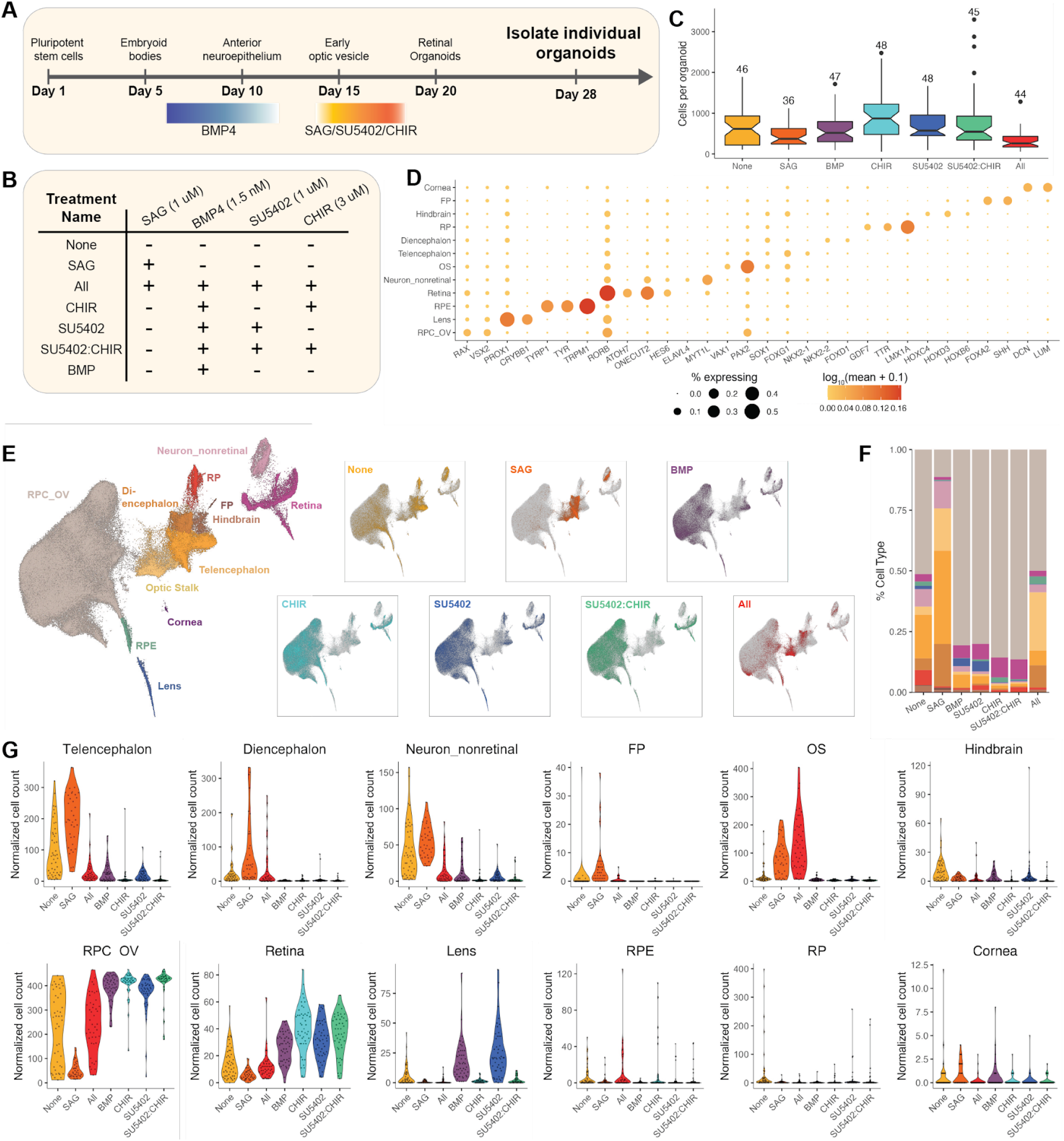
scī-Plex applied to individual organoids after signaling pathway modulation. A) An overview of the retinal organoid differentiation protocol and treatment timing. On Day 6, BMP4 was added to the cultures. BMP4 was diluted in half on Day δ and again on Day 10 before being removed completely on Day 12. SAG, SU5402, and/or CHIR were added on Day 14-18. The organoids were cultured for an additional 10 days before collection. B) A summary of the seven treatments performed in this experiment. Note that the All, BMP, CHIR, SU5402, and SU5402:CHIR treatments include BMP4. **C)** A box plot of the number of cells recovered per organoid for each treatment. The number above the boxplot indicates the number of individual organoids in which > 50 cells were recovered for that treatment. **D)** Dot plot of the genes used to define cell types for D28 organoids. The size of the dot indicates the percent of cells that express the gene of interest. The color indicates the Iog10 mean UMIs per cell. E) Left: UMAP generated by Monocleδ of the recovered cells with cell type annotations from D28 organoids treated with signaling modulators. Right: For each of the UMAPs only the cells from the specified treatment are colored by treatment. RPC_OV: retinal progenitor cell/optic vesicle, RPE: retinal pigmented epithelium, OS: optic stalk, RP: roof plate, FP: floor plate. **F)** A stacked bar plot in which the mean cell type composition of the D28 organoids are displayed for each treatment. **G)** Violin plots of size-factor normalized cell counts for each cell type from D28 organoids. Plots are colored by treatment.

However, we first needed to define cell classes and tabulate their abundances across treatments. Despite the lower UMI counts with the highest throughput method, cell classes were readily defined with clustering followed by the analysis of marker gene expression (**Figures 2D-E, S2H, Table S1**). The largest population of cells was the RPC/optic vesicle (RPC_OV). We also detected cells with characteristics of various retinal neuron precursors. In addition to these retinal cell classes, a large number of non-retinal cells–most of which expressed markers of various parts of the developing brain–were recovered (**Figures 2D-E** and **S2G**). We found that differences in RPC_OV content appeared to vary across treatments (**Figures 2E-G**), most dramatically RPC_OV cells decreased in the sonic hedgehog (SHH) activation (SAG) treatment when compared to the None treatment, and they increased in the BMP, SU5402, CHIR, and SU5402:CHIR treatments (**Figures 2E-G**). Observations of such substantial and sweeping changes could be broadly made, but conclusions about changes in cell class abundance at finer resolution were more difficult to make.

We next aimed to use statistical methods to assess the treatment-dependent changes in cell class abundances to capture both large and small treatment effects. We harnessed the power of our individual organoid data to fit a statistical model using the beta-binomial distribution (Methods)^28, 29^. This method was recently applied to zebrafish embryos where it was highly sensitive in detecting changes even in rare cell classes. To test our model we first wanted to compare our BMP4 and None treatments. Previous reports observed that the addition of BMP4 improves the efficiency of retinal differentiation from stem cells^18^, prompting our group and others to adapt our organoid culturing protocols to routinely add BMP4. Indeed, using our model, the sci-Plex data support the continued addition of BMP4 as we saw that, compared to None, the organoids from treatments that include addition of BMP4, with the exception of All, produced significantly more RPC_OV and retinal cells with most other cell classes decreased (**Figures 2G** and **3A**). We also see that not only did sonic hedgehog activation (SAG) decrease RPC_OV cell counts as posited above, it significantly increased Diencephalon, Telencephalon, Optic Stalk, and Nonretinal neurons compared to the None treatment (**Figure 3A**). Interestingly, Optic Stalk was significantly increased in the All treatment, the only treatment with BMP4 exposure that did not increase RPC_OV/Retina cell counts (**Figure 3A**). Thus, with our well powered experiment and our statistical approach, we were able to interpret cell class level effects in the various treatment groups.

**Figure 3.**
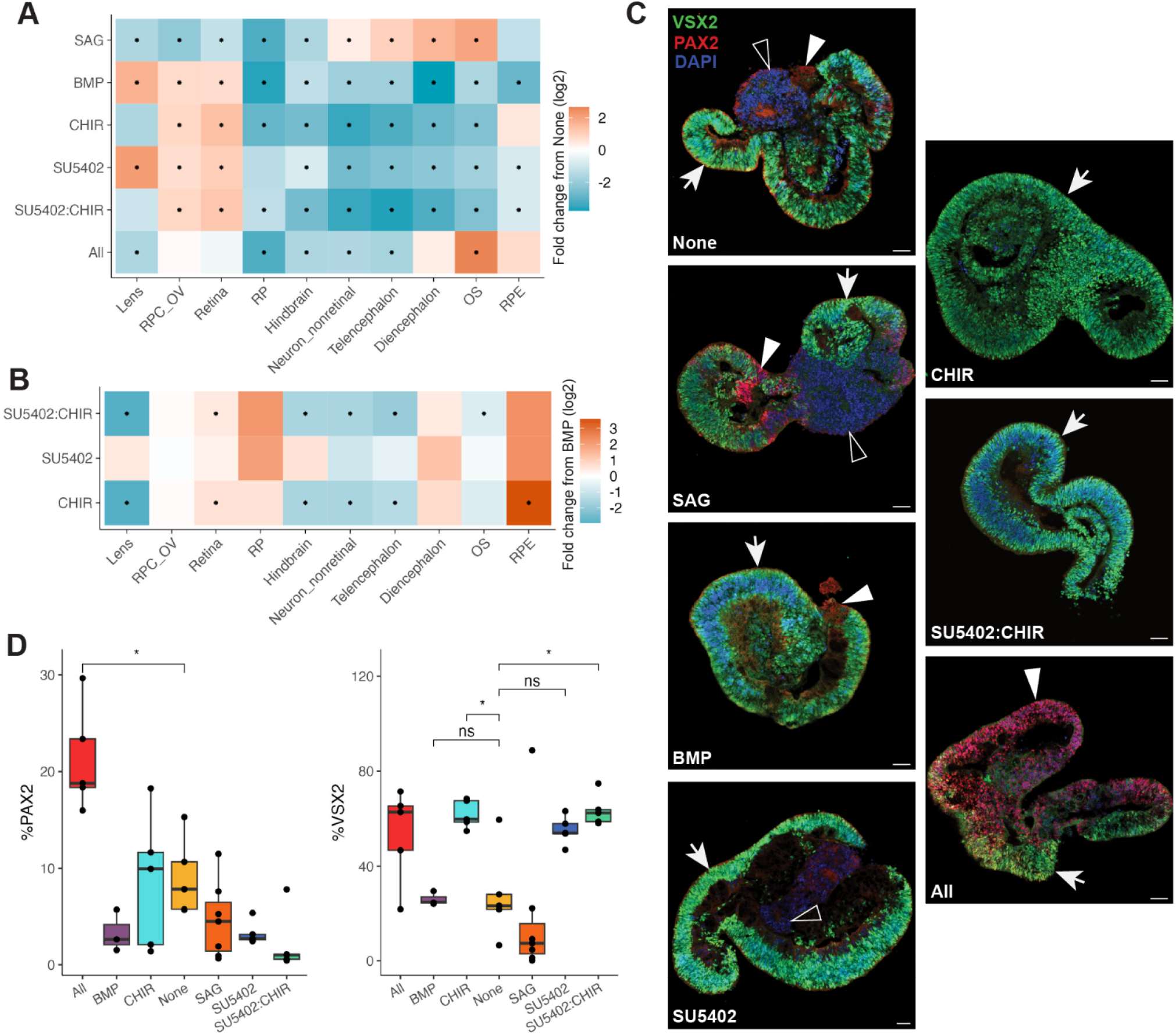
scī-Plex allows the detection of significant changes in cell type abundance. A) Heatmap of fold change in cell type abundance compared to “None" for D28 organoids. A beta-binomial model was fit to the data and used to determine whether there were significant changes in cell type abundances. A* in the center of the box indicates a q-value of less than 0.05 after Benjamini Hochberg correc­tion. RPC_OV: retinal progenitor cell/optic vesicle, RP: roof plate, OS: optic stalk, RPE: retinal pigmented epithelium. B) Heatmap of fold change in cell type abundance compared to “BMP" for D28 organoids. A beta-binomial model was fit to the data and used to determine whether there were significant changes in cell type abundances. A * in the center of the box indicates a q-value of less than 0.05 after Benjamini Hochberg correction. **C)** Immunostaining of D28 organoids with VSX2 (RPCs/Npre, green), PAX2 (OS, red), and DAPI (blue). Scale bar represents 50 µm. Arrow = RPCs/Npre, empty arrowhead = non-optic-stalk (OS), filled arrow head = OS. **D)** Quantification of D28 immunostained organoids. Between 3-7 organoids were stained per condition (All = 5, BMP = 3, CHIR = 5, None = 5, SAG = 7, SU5402 = 5, SU5402:CHIR = 5). Significance was determined by performing t-tests between the treatment of interest and None. Benjamini Hochberg correction was performed to adjust the p-value. * represents a p < 0.05.

After confirming that the BMP treatment was an improvement over None, we next wanted to better investigate whether SU5402, CHIR, or SU5402:CHIR could lead to further advances in the culturing protocol. We trained a new model in which each of the SU5402, CHIR, and SU5402:CHIR treatments were compared to BMP. In our first model, both the BMP and SU5402 treatments led to an increased number of lens cells compared to the None treatment, consistent with the established role of BMPs in lens development (**Figure 3A**)^18, 50, 51^. In contrast, the addition of BMP4 combined with Wnt activation (CHIR, SU5402:CHIR) caused a striking loss of lens cells when compared to BMP (**Figure 3B**). Both treatments with Wnt activation also robustly decreased most other non-retinal cell classes compared to BMP alone (**Figure 3B**). Of the conditions with high RPC_OV cell counts, the CHIR treatment led to the highest relative numbers of RPE cells in the organoids, consistent with prior evidence that Wnt activation steers cells towards the RPE fate both *in vivo* and *in vitro*^50, 52^. Unexpectedly, both the CHIR and the SU5402:CHIR treatments increased neural retinal cell counts (**Figure 3B**). Overall, our results indicated that addition of CHIR99021 to the culturing protocol increased the retinal content of organoids compared to addition of BMP4 only.

We next assayed the subretinal cell class breakdown to see whether any of the treatments tipped the scales towards the production of a subset of retinal cell classes. Using only the RPC_OV and retinal cells, dimensionality reduction was performed and retinal cell classes were determined using marker genes (**Figures S3A-C, Table S2**). CHIR, SU5402, and SU5402:CHIR treatments increased photoreceptor/bipolar cell precursors (PBpre), with CHIR and SU5402:CHIR treatments displaying the most robust increases (**Figures S3D-E**). CHIR and SU5402:CHIR treatments also significantly increased neurogenic precursor (Npre) cells (**Figure S3D-E**). Overall, the results of this experiment show that exposure to CHIR99021 from Day 14-18 reduced non-retinal cell classes, increased retinal cells, and may have specifically enhanced photoreceptor/bipolar cell production at this point in time.

### Cell abundance changes detected by sci-Plex are concordant with immunostaining

To further confirm our observations from sci-Plex, we turned towards immunostaining as a secondary approach. We collected D28 organoids from the same cultures as above and performed immunostaining on cryosections (**Figure 3C**). While immunostaining is a critical tool for the field, it comes with a number of limitations that make it difficult to scale: 1) only a few cell classes can be assayed at a time, 2) it requires prior knowledge of marker genes and existing antibodies to assay the cell classes of interest, and 3) it’s laborious, limiting the number of samples that can be processed. Nevertheless, we observed a significant increase in PAX2+ optic stalk-like regions in the All treatment (**Figure 3D**), consistent with the sci-Plex results. The SU5402, CHIR, and SU5402:CHIR treatments all presented with increased VSX2+ regions (RPCs and retinal neuron precursors), as was detected by sci-Plex, with significant increases in VSX2+ regions in CHIR and SU5402:CHIR treated organoids (**Figure 3D**). However, the immunolabeling results and sci-Plex did not agree when comparing the BMP and None treatments: sci-Plex showed an increase in VSX2+ regions, yet this was not apparent in the cryosectioned organoids. The discrepancy may be due to the modest number of cryosections we had for the BMP treatment compared to the other treatments. sci-Plex captured all of the significant changes observed by immunostaining. However, the reduced number of organoids used for immunostaining and the reduced number of markers we could stain for hindered our ability to find the additional significant changes in cell composition that sci-Plex identified. Overall, we did not detect the same breadth of cell class abundance changes by immunostaining, nevertheless our results were highly concordant between methods.

### Organoid subtypes are identified within treatment groups

In order to quantify heterogeneity in cell proportions across organoids and to assess whether any perturbations reduced it, we looked at the cell class compositions of each individual organoid. Recent innovations have increased the percent of organoids in a given culture that are retinal, but the definition of usable is based on qualitative bright-field images where variation is observable, and the molecular source of variation is not defined^18, 19^(**Figure 4A**). Our individual organoid sci-Plex dataset provided a more quantitative picture of the variation that exists within and across treatments (**Figures 4B-I**) in a population of unselected organoids from a commonly used protocol. Without exposure to any of the signaling modulators (None), organoids ranged from being primarily RPCs to almost completely non-retinal (**Figures 4C**). The organoids from the All treatment were also highly heterogeneous with some individuals containing either RPC, Optic Stalk, or Diencephalon as their most common cell class (**Figure 4D**). BMP treatment markedly reduced the heterogeneity of the organoids, and most of the organoids in the BMP4-treated conditions (BMP, CHIR, SU5402, SU5402:CHIR) were composed of predominantly RPC_OV and retinal cells (**FIgures 4E-H**). This increase in RPC_OV/retinal cells in the individual organoids was particularly apparent in the Wnt activating treatment conditions (CHIR, SU5402:CHIR). Nevertheless, a subset of the organoids demonstrated atypical cell class distributions (**Figures 4E-H**).

**Figure 4.**
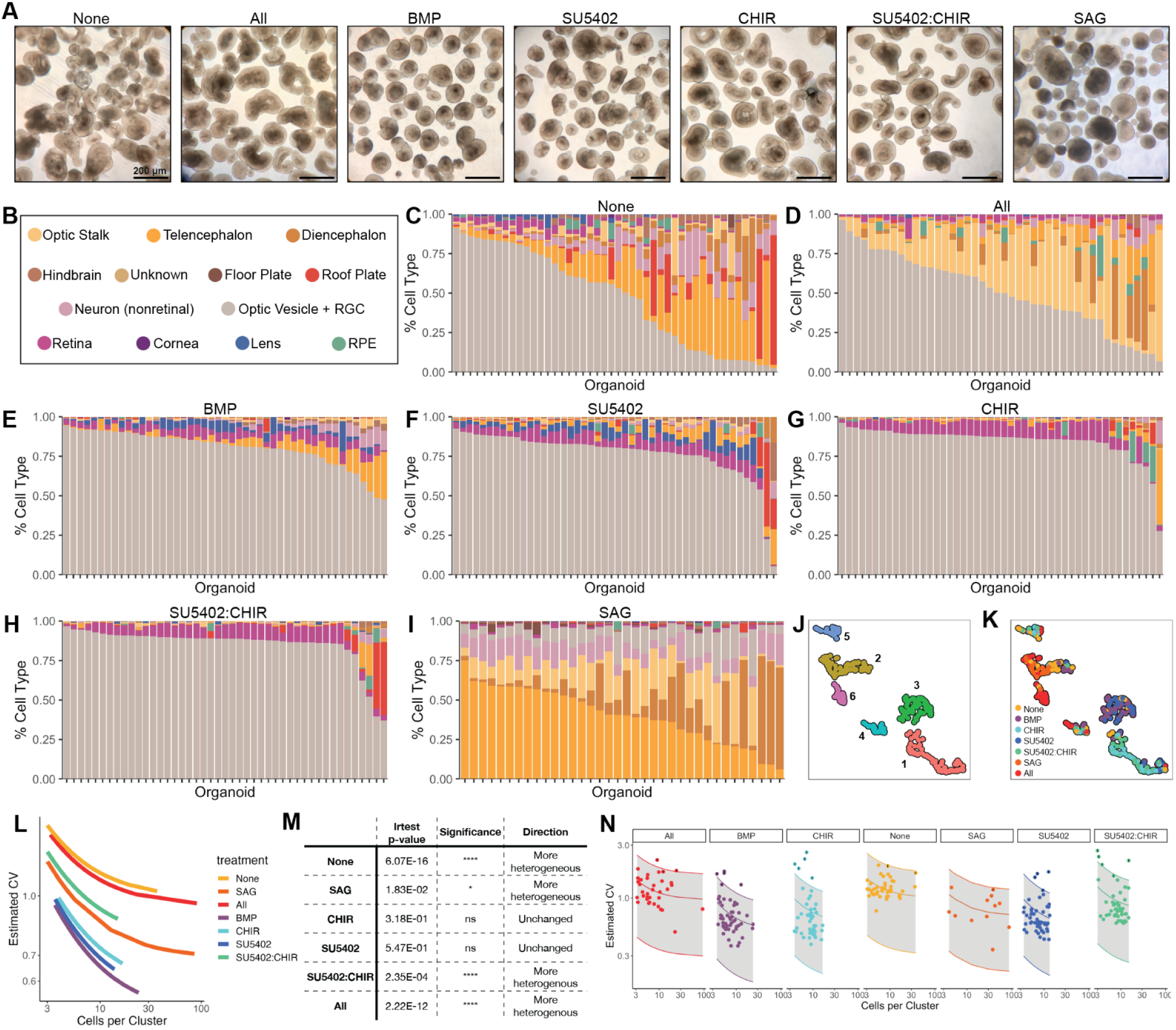
Detection of retinal organoid heterogeneity and organoid subtypes. A) Bright phase images of organoids the day before they were collected for sci-Plex (D27). Scale bars are 200 µM. **B)** Cell type legend for C-l. C) Stacked bar plot in which each bar represents an individual D28 organoid exposed to the None treatment. The bars are colored by the percent a given cell type makes up the composition of the individual organoid. Same as C, but for the **D)** All, **E) BMP, F)** SU5402, **G)** CHIR, **H)** SU5402:CHIR, and **I)** SAG treatments. **J)** A UMAP generated using the size-factor normalized cell type by organoid matrix generated from the D28 individual organoid dataset. Organoids are colored by cluster. **K)** The UMAP from J with the organoids colored by treatment. **L)** The mean and coefficient of variation (CV) were modeled for each cluster and each treatment using a gamma distribution. Displayed are the modeled relationships colored by treatment. **M)** Output of likelihood ratio tests (Irtest) with or without inclusion of the treatment term for D28 organoids. **N)** Model from L faceted by treatment. The shaded area marks the range within 2 standard deviations from the mean. Points represent the individual cluster values.

To define organoids within a culture that are divergent in their cell class compositions we aimed to develop a method to categorize organoids into “archetypes”. We counted the number of cells of each type in each organoid and used the resulting normalized cell class x organoid matrix (see Methods) to create a UMAP in which each of the points represented an individual organoid. We performed clustering and found that organoids of a given treatment fell predominantly into a single archetype (cluster), with the exception of the None treatment which had organoids dispersed across all archetypes (**Figures 4J,K** and **S4A**). It was notable that, across the board, addition of signaling modulators reduced the spread of organoids across archetypes. We were also interested in why a specific secondary archetype may appear for one treatment and not another. For example, Archetype 2, defined by high counts of hindbrain cells and non-retinal neurons, was higher for BMP than it was for the other high retinal cell content treatments (CHIR, SU5402, SU5402:CHIR)(**Figures 4J-K** and **S4B**). All other high retinal content treatments have more organoids from Archetype 5 (high RPE, high roof plate) than BMP alone does (**Figure S4A-B**). These cell classes (i.e. hindbrain, non-retinal neurons, RPE, and roof plate) were not seen as significantly increased compared to BMP (with the exception of RPE in CHIR), yet they were compellingly present in high numbers for a small set of individuals (**Figures 3B** and **4E-H**). Uncovering these cell classes opens the door for designing protocols targeted at eliminating undesirable cell classes and thus increasing organoid homogeneity. Here we have demonstrated the ability to define the typical cellular composition of organoids exposed to a given treatment, and we have detected the manner in which a minority of organoids within a given treatment deviate from the established typical cell class composition.

### Statistical measurements of organoid heterogeneity identify treatments that decrease homogeneity

We next sought to quantify the variability in cell composition and how it increased or decreased with different treatments. To do so, we used a method that accounts for the mean-variance relationship in cell abundances across individuals as previously described^28, 29, 53^. Cells were divided into the 67 clusters we used to define cell classes (**Figure S4C**), and the variance and mean among individuals were calculated for each treatment within each cluster (i.e. the normalized cell counts for the 48 embryos exposed to CHIR treatment were determined for cluster 1 and then the mean and variance were calculated. This was repeated for each of the treatments and then each of the clusters). Because variance scales with the mean, the coefficient of variation (CV) was calculated by taking the ratio of the standard deviation over the mean as a means to normalize the amount of variance such that we can compare the variance in clusters with high cell counts to clusters with low cells counts. As randomness is increased at smaller values, we expect that clusters with fewer cells will have higher CV and clusters with more cells will have lower CV values. After modeling (see Methods), the relationship between the number of cells per cluster and expected CV was plotted (**Figure 4L**). Lines lower on the plot have a lower CV and thus less heterogeneity across clusters. As expected, the CV was low for organoids subjected to the BMP treatment. Comparing BMP to each of the other treatments in a pairwise fashion we used a likelihood ratio test on the models of variance with and without a treatment term in order to assess whether other treatments affected organoid heterogeneity. We found that CHIR and SU5402 treatments did not significantly increase heterogeneity compared to BMP, but all other treatments did (**Figure 4M**). Our modeling also enabled us to identify specific clusters of cells that have a higher CV than statistically expected (**Figures 4N** and **S4D-J**). This included most commonly the RPE and roof plate cell clusters, followed by other non-retinal cell class clusters (**Figures S4D-K**). This analysis used a statistical method in a novel manner to find that the additional activation of Wnt or inhibition of FGF signaling does not change a culture’s homogeneity compared to treatment with BMP4 alone but that all other treatments increase heterogeneity.

### Early modulation of signaling pathways affects long-term cellular composition

At D28, the organoids are mainly progenitors. To see whether the effects of signaling modulation carry forward as the cells differentiate into more mature fates, we performed sci-Plex on 96 individual organoids cultured for 63 days (BMP, CHIR, or SU5402:CHIR treatments). After filtering, 194,335 hashed cells were recovered (median UMIs: 341, median features: 298) with a median of 1841.5 cells per organoid across 96 organoids (**Figures S5A-G**). Cell classes were defined using marker genes (**Figures 5A** and **S5H**). Although retina progenitors and neurons made up the majority of the cells at this stage, cells with brain characteristics persisted. Consistent with the D28 organoids, Wnt activation (CHIR, SU5402:CHIR) led to significant decreases in mean cell counts for non-retinal/cortical neurons and undefined cells when compared to BMP4 alone (**Figures 5B-C**). In addition, there was a significant increase in RPE cells upon FGF inhibition and Wnt activation (SU5402:CHIR)(**Figures 5B-C**). At D63, some retinal neuron cell classes, including PBpre, HApre, RPCprolif, and RPCproneural, were broadly increased in the CHIR and SU5402:CHIR treatments, when compared to the BMP condition, but not to the level of significance (**Figures 5B** and **S5I**). Plotting the individual organoid cell composition (**Figures 5D**) also highlighted the fact that Wnt activation (CHIR) reduced the number of organoids containing high representation of cortical neurons, when compared with BMP treatment. However, overall heterogeneity was not significantly different in any of the tested treatments (**Figures S5K** and **Table S3)**, and non-retinal cell classes tended to be more variable than retinal cell classes (**Figure S5J,L**). Immunolabeling of cryosections of organoids from the same batch used for the sci-Plex analyses confirmed the representation of non-retinal cells in these organoids (**Figure 5E**). Taken together, this suggests that the early modulation of signaling pathways establishes durable shifts in retinal cell class representation.

**Figure 5.**
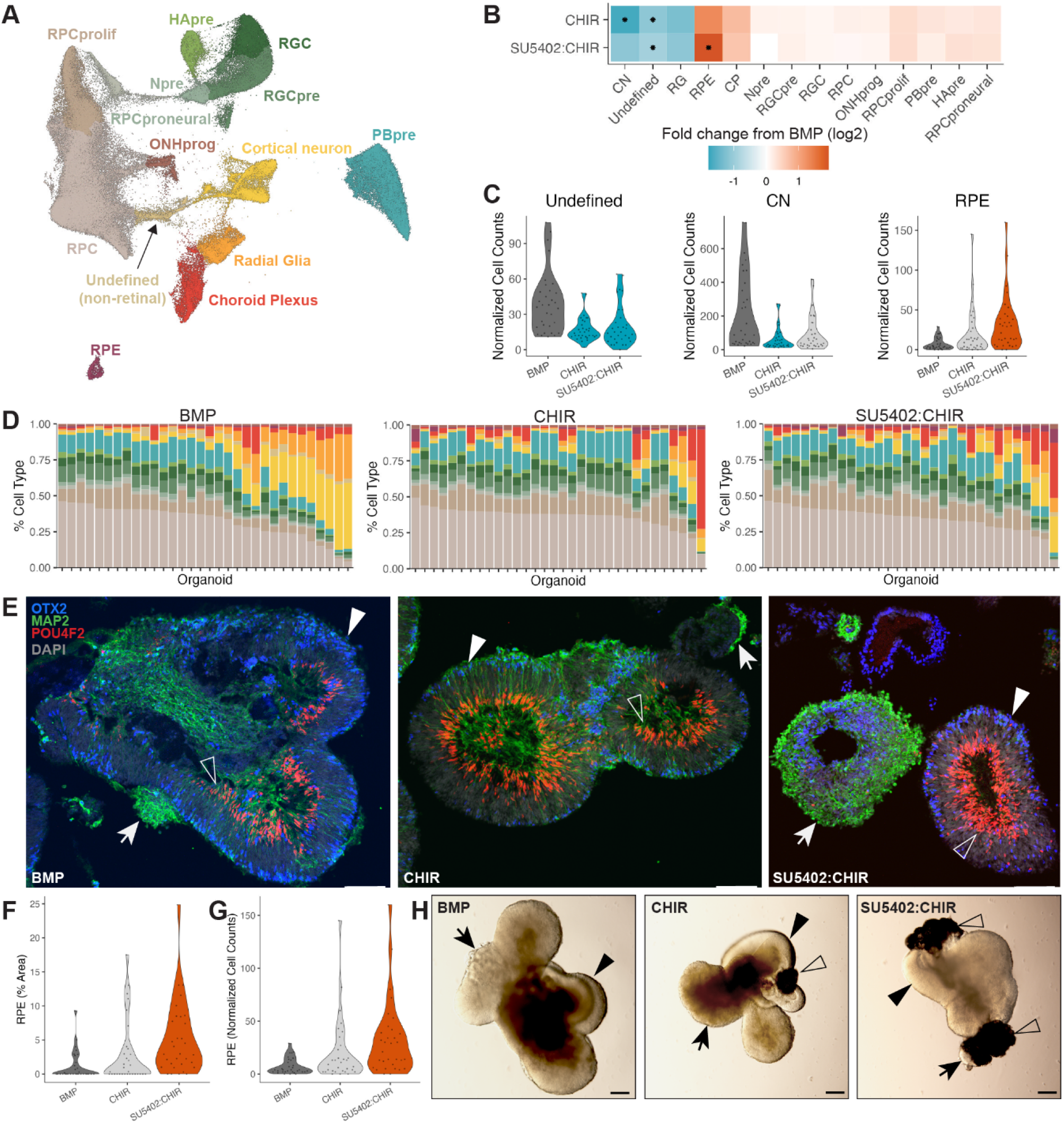
D63 organoids demonstrate continued perturbation by signaling modulation. **A)** UMAP from Mono­cles with cell type annotations of all cells recovered from individually hashed D63 organoids exposed to BMP. CHIR, and SU5402:CHIR treatments. RPC: retinal progenitor cell, RPCprolif: proliferating RPC, RPCproneural: proneural RPC, Npre: retinal neuron precursor, HApre: horizontal/amacrine precursor, RGC: retinal ganglion cell, RGCpre: RGC precursor, PBpre: photoreceptor/bipolar precursor, ONHprog: optic nerve head progenitor, RPE: retinal pigmented epithelium, CN: cortinal neuron, RG: radial glia, CP: choroid plexus. B) Heatmap of fold change in cell type abundance compared to BMP. A * in the center of the box indicates a q-value of less than 0.05 after Benjamini Hochberg correction. C) Violin plots of size-factor normalized cell counts for D63 organoids. Dark gray = the treatment that was used for fold-change calculations in B, light gray = no significant change in cell type abundance, teal = decrease in cell type abundance, orange = increase in cell type abundance. D) Stacked bar plots in which each bar represents an individual organoid exposed to the BMP, CHIR, or SU5402:CHIR treatment. The bars are colored by the percent a given cell type makes up the composition of the individual organoid. Right: cell type color legend. E) Immunostaining of cryosectioned D63 organoids with OTX2 (photoreceptors/RPE/CP, blue), MAP2 (retinal and cortical neurons, green), POU4F2 (RGC, red), and DAPI (gray). Arrow = Non-retinal neurons, filled arrow head = Otx2+ retinal tissue, empty arrow head = Pou4f2+ retinal tissue. Scale bar represents 100 µm. **F)** Violin plots of % Area measurements for RPE as calculated from bright phase microscopy images of D62 organoids. Plots are colored by significance as in C. Violin plots of size-factor normalized RPE cell counts for D63 organoids. Plots are colored by significance as in C. Representative brightfield images of D62 organoids from each group quantified in G-H. Arrow = non-retinal tissue, filled arrow head = retinal tissue, empty arrow head = RPE. Scale bar represents 200 µm.

Lastly, we compared the cell counts obtained from sci-Plex to measurements of organoid composition obtained by bright-field microscopy. Each organoid was imaged in a well of a 96-well plate such that the imaged organoid could be matched to a specific hash barcode. Areas of predicted retina, RPE, and non-retinal tissue were quantified and compared to the sci-Plex cell counts (**Figures 5F-H** and **S5M-N**). Despite differences in the modalities, general trends were conserved between the imaging and sci-Plex datasets. Notably, RPE was significantly increased in SU5402:CHIR by both sci-Plex and bright-field microscopy (**Figures 5B,F**). The cell count and area-based measurements from each individual also correlated although sci-Plex detected a higher retinal composition and the bright-field images detected higher RPE and non-retinal composition (**Figure S5O**). Overall, we have shown agreement between two highly divergent methods. However, our analysis by bright-field microscopy was limited to a quantification based on broad cell class domains using 2D images of a 3D object. The ability to quantify all cell classes across hundreds of individual organoids in a manner that agrees with and surpasses the sensitivity and breadth of the next best methods sets sci-Plex apart as a critical new method for the study of individual organoid cell class composition across treatment conditions.

## Discussion

Plate-based screening assays that produce cells with desired phenotypes are widely used to identify treatments for a range of diseases. With the increasing utility of organoid culturing systems, it follows that demand for plate-based methods to analyze a perturbation’s effect on organoids has grown. However, current high-throughput techniques don’t have the resolution to capture cell-type specific information across all cell classes simultaneously. Traditional single-cell sequencing technologies provide a high depth of cell type-specific information, but they are limited in scale. With an increased throughput, the original sci-Plex demonstrated its utility for drug screening in cancer cell lines - here we show that it also promises to help us perform screening in culturing systems with complex cell class communities. We successfully use sci-Plex to capture a diverse array of cell classes across 410 individual retinal organoids treated with BMP, Wnt, Shh, and/or FGF pathway modulators, and it enabled us to identify CHIR99021 as a small molecule that, when added to the existing culturing protocol during the third week, reliably decreases non-retinal cell classes without increasing organoid-to-organoid heterogeneity.

By using a high number of individual organoids in each experiment, previously intractable analyses are now possible. We identify significant changes in cell class abundance with a statistically principled methodology. We are also able to, for the first time, comprehensively measure the extensive heterogeneity of retinal organoids, a known but poorly quantified feature of the current retinal organoid culturing protocols. Recent publications emphasize the utility of using single-cell sequencing to investigate individual organoids and organoid heterogeneity^30, 31^. However, no other study has profiled cell class composition in comparable sample sizes across multiple treatments (32-48 individuals per treatment). Our work represents the most complete investigation of cell class composition and heterogeneity in any organoid system to date.

With our methodologies, we find that cell class abundance changes expected from the literature are readily detected. BMP treatment increased retinal and lens character in organoids^18, 50, 51^, Wnt activation in the optic vesicle promoted RPE fate determination ^48^, and the shift towards organoids with greater brain identity upon SAG treatment reflects the role of SHH signaling in defining the ventral forebrain^54, 55^. All together, these results agree with the biology of eye development as well as recent observations in retinal organoid culturing techniques.

Additionally, we argue that non-retinal tissues in retinal organoids merit greater attention than they have previously commanded. Tissue that appears non-retinal by bright-field microscopy is routinely dissected away to improve the homogeneity of the cultures. However, when adding all four of our chosen signaling modulators in the All treatment, we generate organoids with optic stalk-like cells, a cell type that is of clinical relevance but for which an established 3D culturing system does not exist^56^. Optic stalk cells would have been discarded as non-retinal by conventional bright-field microscopy; however they are captured in our study by the unbiased approach we used to survey the entire composition of organoids at scale. Further use of the technology in the context of retinal development and its ability to be applied in other developmental contexts has the potential to uncover novel culturing systems for additional disease-relevant cell types that may form unexpectedly or be difficult to detect by traditional methods.

Another advantage of detecting all cell classes within an organoid, specifically when it can be performed across individuals, is the ability to clarify the sources underlying organoid heterogeneity. We show that it’s possible to separate individual organoids into subtypes based on their cell class composition. With CHIR treatment, > 75% of organoids fall into a single subtype defined by high RPC and retinal cell counts, but a secondary subtype defined by high RPE and Roof Plate content persists. What is driving this heterogeneity? Can we apply additional developmental signals that specifically regulate roof plate development in order to reduce heterogeneity^57^? Could we layer in additional signaling modulators to more uniformly induce or repress the RPE cell fate choice?

When we performed the analysis of organoid-to-organoid heterogeneity, we noticed that the None and All treatments were intriguingly the most variable. That the addition of signaling molecules leads to the canalization of cells towards Brain (SAG) or Retinal (BMP) fates with increased homogeneity indicates that under the None culturing condition the cells are not robustly restricted in their fate decisions. To the contrary, in the All treatment, four different signaling molecules are added, and individual variation remains high. We predict that the modulation of so many signaling pathways could lead to competition and interference that destabilize differentiation trajectories. The *in vitro* recapitulation of development is a complex dance that requires a balance of signals at the right time to produce organoids of desired composition. We show here that both the input of not enough and too many signals detrimentally affects reproducibility.

Lastly, our most surprising result is the CHIR-dependent increase in neural retina at D28 and the decrease in non-retinal neurons through D63. Active canonical Wnt-signaling is associated with decreased telencephalic identity and increased diencephalic identity, but the role of Wnt in determining the eye field is less well-understood^58, 59^. Was the increase in retinal cells at D28 a result of the loss of non-retinal cells or does Wnt promote the retinal fate? We look forward to further teasing apart the regulatory mechanisms driving these changes in cell identity.

Ultimately, the methodological advancement of sci-Plex and our analytical pipeline facilitates the expansion of perturbation studies in organoids. While we tested 7 conditions across 410 individuals, sci-Plex can readily be expanded to thousands of individuals across as many conditions as are desired. We imagine this will lead to continued improvements to organoid culturing protocols, as well as an expansion in the use of organoids in chemical screening platforms, and in understanding the mechanisms of human development. Although there are many applications for sci-Plex, we envision its utility in the near future to identify ganglion cell survival molecules and to screen for molecules that aid in cone survival as potential treatments for macular degeneration. Outside the retina, this technology holds promise for use with the constantly expanding number of 3D culturing systems for which there is a demand to increase the throughput of samples to be sequenced at single-cell resolution.

### Limitations of Study

We designed this study in favor of a high number of individual organoids in order to sample the organoid-to-organoid variation in an unprecedented fashion. Our choice revealed that indeed high counts of individual organoids improve the ability to detect changes in cell class abundances, however we did not test all possible culturing conditions of interest including but not limited to treatment of organoids with other small molecules or the testing of different organoid culturing protocols. Additionally, our analyses focused primarily on cell class abundances instead of gene expression for two reasons: 1) sci-RNA-seq3 returns low UMIs per cell, although newer techniques have improved upon this^60^. The recovery is sufficient for broad cell class assignment, but more UMIs per cell may be able to further segregate similar cell classes and increase the detection of DEGs. 2) The treatments were performed weeks before the organoids were collected. We wanted to allow the organoids time to specify cell fates after the treatments were administered. Thus, the effects on gene expression caused at the time of treatment were not captured. This prevents us from establishing a more mechanistic understanding of the regulatory events that lead to the long-term changes in cell abundance that we observe. Lastly, in our experiments that use hashing with sci-RNA-seq3 our hash rates were ∼50%. This means that we are sequencing many more cells than can reliably be assigned to an individual and a shortcoming we are currently working to minimize. Furthermore, the original sci-RNA-seq3 protocol has a low cell recovery rate, so this protocol isn’t ideal for rare cell types.

## Acknowledgements

We’d like to thank the members of the Trapnell and Reh lab for their valuable comments and discussions. We thank Choli Lee for assistance in flow sorting, the Beliveau lab for use of their Keyence BZ-X810, and the Brotman Baty Institute Advanced Technology Lab for support with the data processing pipeline. Funding: this work was supported by the National Institutes of Health (1R01HG010632 to C.T.; R01EY021482-12 to T.A.R; and 1F32EY032331 to A.T.), the Paul G. Allen Frontiers Foundation (Allen Discovery Center grant 12357 to C.T.), the Chan Zuckerberg Initiative (CZF2019-002442 to C.T.), the Foundation Fighting Blindness (TA-RM-0620-0788-UWA to T.A.R.), the Damon Runyon Cancer Research Foundation (DRG-# 32-20 to K.C.E.), and the HHMI Hannah Grey Fellowship to K.C.E. Illustrations were created with BioRender.com.

## Author Contributions

Conceptualization, AT, AS, KE, TR

Methodology, AT, CT

Formal Analysis, AT, KE, AS

Investigation, AT, AS, KE, DH, SC

Writing-Original Draft, AT

Writing-Review & Editing, AT, KE, CT, TR

Visualization, AT, AS, KE

Supervision, CT, TR

Funding Acquisition CT, TR

## Methods

### Resource availability

#### Lead contact

Further information and requests for resources and reagents should be directed to and will be fulfilled by the lead contact, Thomas A. Reh.

#### Materials availability

No unique materials were generated by this study.

### Stem cell and retinal organoid culture

Parent cell lines used were the human embryonic stem cell lines (hESCs) H9 (WiCell, Madison, WI, https://www.wicell.org). H9 hESCs contain an *NRL*^+/eGFP^ reporter ^51^contain. All organoids of the same age were from the same batch, but each age was from a different starting batch of organoids. The time matched sci-Plex and immunostaining experiments used organoids from the same batch.

| Experiment | Age | Line | Organoids Used |
| --- | --- | --- | --- |
| 10x match | D78 | H9 | 5 pooled |
| 10x match | D185 | H9 | 5 pooled |
| 10x match | D29 | H9 | 10 pooled |
| D28 | D28 | H9 | 314 individuals |
| D63 | D63 | H9 | 96 individuals |

hESCs were maintained by dissociation and propagation in StemFlex media (ThermoFisher, A3349401) on Matrigel (Corning, CLS354277) coated plates at 37 degrees C in 5% CO_2_. hESC colonies were passaged by dissociation with ReLeSR (StemCell Technologies, 100-0484). Retinal organoids were grown as described in^18^ with minor alterations mentioned herein. The cell culture medias used were Neural induction media, or NIM: 484.5mL DMEM/F12 (Life Technologies, Catalog #11330-057), 5mL N2 supplement (Life Technologies, Catalog #17502048), 5mL MEM NEAA (Life Technologies, Catalog# 11140050), 5mL penicillin-streptomycin (Life Technologies, Catalog# 15240062). Retinal differentiation media, or RDM: 240mL DMEM/F12 (Life Technologies, Catalog #11330-057), 240mL DMEM (Life Technologies, Catalog #12430062), 10mL B27 supplement (Life Technologies, Catalog #17504001), 5mL MEM NEAA (Life Technologies, Catalog #11140050), 5mL penicillin-streptomycin (Life Technologies, Catalog #15240062) + 1%, 3%, or 5% FBS (Corning, Catalog #35-011-CV) based on protocol needs.

In brief, confluent stem cell colonies were lifted from the plate using dispase (2mg/mL, Life Technologies, Catalog #17105041) for 5-10 min., then forcefully removed by pipetting 2 mL/well of DMEM (Life Technologies, Catalog #12430062) directly onto cells using a 1 mL pipette. Colonies were removed from the 6 well plates and allowed to settle to the bottom of a 15 mL collection tube by gravity. The supernatant was removed, and colonies were transferred to a T25 tissue culture flask in a 3:1 StemFlex:NIM mix for a total of 10 mL per flask. Full media change was performed the following day (Day 1) with 10 mL of 1:2 StemFlex:NIM. Day 2 a full media change was performed and replaced with 10 mL of NIM. Day 3 a full media change was performed and replaced with 10 mL of NIM. Day 5 a full media change was performed and replaced with 10 mL of NIM. Day 7 a full media change was performed and replaced with 10 mL of NIM, and BMP4 BMP4 (1.5 nM, R&D Systems, 314-BP-050) was added to the media to a final concentration of 1.5nM. Day 8 EBs were then evenly distributed between the 6 wells of a 6 well plate. The media was not changed, and EBs were moved and cultured in the BMP4 containing NIM media they were in. 200 uL of FBS was added to allow for EBs to stick to the bottom of the plate. Days 10-20, every other day, a full media change was performed with fresh NIM. Days 14 and 16 a full media change was performed with NIM, and the signaling factor modulators CHIR99021 (3 uM, Cedarlane Laboratories, 04-0004-02), SU5402 (1 uM, Cayman Chemical Co, 13182), and/or SAG (1 uM, Calbiochem, 566660) were added depending on the treatment group. For the None treatment, DMSO was added at the same volume and time in which SU5402 was added. EBs were then manually lifted from the plates on day 20 or 21 by forcefully pipetting 2 mL/well of DMEM directly onto EBs using a 1 mL pipette. EBs were removed from the 6 well plates and allowed to settle to the bottom of a 15 mL collection tube by gravity. From this point on we consider the cells retinal organoids. The supernatant was removed, then organoids were resuspended in RDM with 1% FBS. Two days later media was exchanged with RDM +3% FBS, and two days after that with RDM + 5% FBS. From this point on, cells were maintained with a full media change of RDM +5% FBS every two to three days.

### Single cell dissociation

Organoids were placed into 96-well v-bottom plates. For the 10x comparison, 5-10 organoids were added to each well. In the D28 and D63 experiments, a single organoid was added to each well. Media was removed, the organoids were washed in DPBS (no calcium, no magnesium) (ThermoFisher, 14190144). For the D28 and D63 organoids, 100 uL of accutase (Sigma, A6964-100ML) was added to the cultures. The cultures were incubated at 37 °C on a nutator for 15 minutes. After 15 minutes, organoids were pipetted up and down as needed (every 5-10 minutes). The extent of dissociation was determined by bright-field microscopy. When clusters of cells could no longer be seen, the accutase was deactivated by the addition of 20 ul of FBS (Corning, 35-011-CV). This was done on an organoid-by-organoid basis as some dissociate faster than others. The dissociation was complete by 30-45 minutes after accutase addition. For the D78 and D185 organoids, the dissociation was performed with 100 uL of papain (Worthington Biochemical, LK003150). The dissociation was performed as above and was completed within 15-20 minutes. 100 uL of Ovomucoid (Worthington Biochemical, LK003150) was used to deactivate the reaction. After deactivation of accutase or papain cells were spun down at 700 g for 3 min, then resuspended in a single cell suspension in RDM media ^18^ for sci-Plex processing.

### Immunostaining

Organoids were fixed in 4% paraformaldehyde, 5% sucrose for 30-45 minutes followed by 3 washes in 1X PBS. All samples were then transitioned into 10%, then 20% and then 30% solutions of sucrose for at least 30 min each solution. Organoids were then embedded in OCT and cryosectioned at 14-16 mm. For immunostaining, slides were washed three times in 1X PBS, then blocked in blocking buffer (0.5%Triton and 10% horse serum) for 20-60 min, followed by addition of primary antibody diluted in blocking buffer and incubation at 4C overnight. The following day, slides were washed 3 times in 1X PBS, then the appropriate 488/568/647 fluorophore-conjugated secondary antibody, diluted in blocking buffer, was added and incubated for 1 hour at room temperature. Slides were mounted using Fluromount-G (SouthernBiotech, 0100-01). Details of the antibodies used in this study are provided below.

| Antibody | Source | Catalog # | Dilution |
| --- | --- | --- | --- |
| POU4F2 | Santa Cruz Biotechnology | SC-6026 | 1:300 |
| PAX2 | BioLegend | 901001 | 1:300 |
| VSX2 | Exalpha Biologicals | X1180P | 1:300 |
| VSX2 | SantaCruz | Sc-365519 | 1:300 |
| MAP2 | Sigma | M9942 | 1:200 |
| OTX2 | ABCAM | 21990 | 1:300 |
| ELAVL3/4 | Invitrogen | A-21271 | 1:100 |
| RCVRN | Chemicon | AB5585 | 1:200 |

### Microscopy

For brightfield images, single organoids were placed into each well of a round-bottom 96-well plate and imaged at 4x magnification with a Keyence BZ-X810 microscope. Group bright-field images were obtained at 4x magnification using the Zeiss Axio Observer D1. Confocal images were obtained using the Zeiss LSM 990. Image processing was done in ImageJ^61^. Area measurements were obtained by setting a standardized threshold per channel.

### 10x library preparation

Dissociated cells were subjected to the standard 10x workflow for Single Cell 3’ v3.1 Dual Index Gene Expression kit (Dual Index Kit TT Set A 96 rxns, 10x Genomics, 1000215).

### sci-Plex (hashing)

Hashing of organoids uses a protocol adapted from the original sci-Plex publication ^27^. Single-cell suspensions in 96-well v-bottom plates were centrifuged at 600 x g for 5 minutes. This step and all following centrifugation steps were performed in a swinging bucket centrifuge that was chilled to 4°C. Media was removed by aspiration. Cells were washed with 200 uL 1 x dPBS (no calcium, no magnesium) in each well. After centrifugation of the v-bottom plate, the dPBS was removed from wells by aspiration, and 50 uL of CLB+hash solution (45 uL of Cold Lysis Buffer – 10mM Tris/HCl pH 7.4, 10mM NaCl, 3mM MgCl2, 0.1% IGEPAL, 1% (v/v, Sigma-Aldrich, I8896), SuperaseIn RNase Inhibitor (20 U/µL, Ambion, AM2694), 1% (v/v) BSA (20 mg/ml, NEB, B9000S) + 5μL of hash oligo (10 uM, IDT)) was added to the well. The cells were mixed with the CLB+hash by pipetting up and down for 5-10 strokes. The plate was allowed to sit on ice for 3 minutes. To fix the hashes to the nuclei, 200 uL of fixation buffer (5% paraformaldehyde (EMS, cat. no. 50-980-493), 1.25x dPBS) was mixed with the nuclei by pipetting up and down for 10 strokes. The nuclei were fixed on ice for 15 minutes and then all wells of a plate were pooled into a single 50 mL conical tube. The hashed nuclei were centrifuged for 5 minutes at 800xg. Pellets were resuspended in 1mL of NSB (Nuclei Buffer + SuperaseIn + BSA – 10mM Tris/HCl pH 7.4, 10mM NaCl, 3mM MgCl2, 1% (v/v) BSA, 1% (v/v) SuperaseIn RNase Inhibitor) and all 50 mL tubes were pooled into a single 50 mL conical. Nuclei were centrifuged for 5 minutes at 800 x g and resuspended again in 1 mL of cold NSB. Nuclei were transferred to a 1.5 mL LoBind microcentrifuge tube (Eppendorf, Z666491). The number of nuclei recovered was counted using a hemocytometer. In order to continue centrifuging in a swinging bucket centrifuge, the 1.5 mL tube was placed within a 15 mL conical. The nuclei were centrifuged one final time for 5 minutes at 800 x g and they were resuspended in 500 uL of NSB. Samples were flash frozen in liquid nitrogen and stored at -80C.

### sci-RNA-seq

The sci-RNA-seq experiment described in Figure 1 was prepared by the original sci-RNA-seq protocol^32^ with minor modifications. Briefly, fixed nuclei were thawed and centrifuged at 800 x g for 5 minutes. Nuclei were permeabilized with 500 uL NSB + 0.2% Triton X-100 (ThermoFisher, A16046.AP) on ice for 3 minutes. Nuclei were centrifuged for 10 minutes at 800 x g and resuspended in 400 uL NSB. A final centrifugation step was performed at 800 x g for 10 minutes. The nuclei were diluted to ∼300 nuclei per uL across 2 x twin.tec™ 96 Well LoBind PCR Plates (Eppendorf, 0030129512). The reverse transcription reaction was set up as in Cao et al., and the RT reaction was carried out using an increasing temperature gradient (4°C 2 min, 10°C 2 min, 20°C 2 min, 30°C 2 min, 40°C 2 min, 50°C 2 min, 55°C 15 min) without the addition of any reagents to stop the reaction upon completion. All nuclei were pooled and stained with DAPI (4′,6-diamidino-2-phenylindole, 3 uM final) (Invitrogen, D1306). Using a FACSAria III cell sorter (BD Biosciences), 50 nuclei were sorted into each well of 4 twin.tec™ 96 Well LoBind PCR Plates that had 5 uL of EB buffer (Qiagen, 19086), 0.5 uL of 5 x mRNA Second Strand Synthesis buffer (New England Biolabs, E6111L), and 0.25 uL of mRNA Second Strand Synthesis enzyme (New England Biolabs, E6111L) in each well. The plates were incubated at 16°C for 3 hours. To each well, 5.75 uL of Tagmentation mix (1.1 uL N7 loaded custom TDE1 enzyme (MacroLab, UC Berkeley) per 632.5 uL 2xTD buffer (20 mM Tris-HCl pH 7.6, 10 mM MgCl_2_, 20% v/v Dimethyl Formamide (Sigma-Aldrich, 227056-100ML)) was added and mixed by pipetting 10 x. Tagmentation was carried out at 55C for 5 minutes before being stopped by the addition of 12 uL of DNA binding buffer (Zymo Research, D4004-1-L) for 5 minutes at room temperature. Ampure-based bead purification, PCR, and final library purification was performed as published previously ^32^. The library was visualized using a D1000 Screen Tape (Agilent Technologies, 5067-5583).

Both of the individual organoid datasets were prepared following the sci-RNA-seq3 protocol from Srivatsan et al. with modifications noted below. As above, all centrifugation speeds were performed at 800 x g. Fixed nuclei were thawed and centrifuged for 5 minutes. Nuclei were permeabilized with 500 uL NSB + 0.2% Triton X-100 on ice for 3 minutes. Nuclei were centrifuged for 10 minutes and resuspended in 400 uL NSB to wash away the Triton. The nuclei were sonicated with a Bioruptor Plus (diagenode) sonication device on low power for 12 seconds. A final centrifugation step was then performed for 10 minutes. Nuclei were suspended to a concentration of 1000-2500 nuclei/uL with 22,000-55,000 nuclei added to each well of 2 x twin.tec™ 96 Well LoBind PCR Plates. From here only minor modifications from the Srivatsan protocol were performed. Of note, before second strand synthesis 2500-3000 nuclei were distributed into each well, during tagmentation the 10 µL of tagmentation mix was composed of 0.05 µL of a custom TDE1 enzyme in 9.95µL of 2 x TD buffer, and when performing the well-based Ampure bead purification the volume of beads was increased to 60 uL per well. PCR was performed for 13 cycles and libraries were purified with column-based DNA purification (Zymo Research, D4014) followed by 0.8x Ampure bead cleanup. Libraries were visualized using D1000 or HS D1000 Screen Tapes.

In-depth protocols can be found on protocols.io under the following titles.

<u>2- level:</u> Single cell RNA sequencing library preparation (2-level sci-RNA-seq)
<u>3- level:</u> sci-RNA-seq3

### Processing of single cell RNA sequencing data

#### 10x

The 10x libraries from D28, D78, and D185 organoids were sequenced on an Illumina NextSeq 2000 and a NextSeq 550 (Read 1: 28, Index 1: 10, Index 2: 10, Read 2: 75) and processed by CellRanger (3.1.0)^62^ using the default settings. The output was imported into Monocle3 (1.0.0)^33^, and data from the three runs were combined into a single object with the corresponding sci-Plex dataset.

#### sci-Plex (10x comparison)

The sci-Plex library was sequenced on an Illumina Nextseq550 (High Output 75 cycle kit) with 18 cycles for Read1, 10 cycles for each index, and 52 cycles for Read2. The reads were processed using a pipeline developed by the Brotman Baty Institute (BBI). This BBI pipeline is available at the bbi-lab github page under the bbi-dmux and bbi-sci repositories (https://github.com/bbi-lab). The output was imported into Monocle3, and custom code was used to incorporate the hash information outputted by the pipeline into the Monocle3 cds object. Cells with fewer than 1000 UMIs and > 20,000 UMIs were discarded. Hashing success was determined by the ratio of the most prevalent hash barcode for a cell compared to the second most prevalent hash barcode. If the top_to_second_best_ratio was > 3 the cell was considered hashed. All cells below that cutoff were discarded. The filtered sci-Plex data was combined with the 10x data. The combined object was converted into a Seurat object for CCA integration following the “Introduction to scRNA-seq integration” vignette^34, 63, 64^. The integration anchors were found using 100 dimensions and 100 anchor features. A 3D UMAP was constructed from 50 PCA dimensions with a mindist of 0.2 and 50 nearest neighbors. The UMAP coordinates outputted from the integration were transferred back to Monocle3 from which the remainder of the analysis and plotting was performed. Clusters were determined using cluster_cells() in Monocle3 using the default assigned resolution. Common marker genes for retinal cell classes compiled from previous single cell and bulk transcriptomic studies of retinal development in fetal human (from our group and others) were then plotted on to the UMAP to determine the clusters with which they were most associated (Table S2). For some clusters, common retinal markers were not sufficient to assign cell classes. In those cases, genes with high specificity to that cluster were determined using custom code that will be made public upon publication. These genes were then used to aid in a cell class determination (Table S2).

#### sci-Plex (D28 individual organoids)

The sci-Plex library was sequenced on an Illumina NextSeq 2000 (P2 100 cycle kit) with 35 cycles for Read 1, 10 cycles for each index, and 70 cycles for Read2. Reads were processed using the BBI pipeline as described above. The output was imported into Monocle3. A preliminary filtering was performed for cells with > 100 UMIs, less than 20,000 UMIs, and < 15% mitochondrial reads. Cells with a top_to_second_best_ratio < 3 were also removed from the analysis. Size factors were estimated, the data was preprocessed with 40 PCA dimensions, correction was performed to account for the number of UMIs per cell. UMAPs were projected in 2 dimensions with a mindist of 0.1 and 15 nearest neighbors. Cells were clustered with a resolution of 5x10^-5^. cell classes were assigned using well-established marker genes (Table S2). For the analysis of individual organoids, only organoids with at least 50 cells were used.

#### sci-Plex (D63 individual organoids)

The sci-Plex library was sequenced on a Nextseq 2000 (P2 100 cycle kit) with 35 cycles for Read 1, 10 cycles for each index, and 70 cycles for Read2. Reads were processed using the BBI pipeline as described above. The output was imported into Monocle3. A preliminary filtering was performed for cells with > 150 UMIs, less than 20,000 UMIs, and < 15% mitochondrial reads. Cells with a top_to_second_best_ratio < 2.5 were also removed from the analysis. Size factors were estimated, the data was preprocessed with 30 PCA dimensions, correction was performed to account for the number of UMIs per cell. UMAPs were projected in 2 dimensions with a mindist of 0.1 and 50 nearest neighbors. Cells were clustered with a resolution of 5x10^-5^. cell classes were assigned using well-established marker genes (Table S2). Only organoids with at least 50 cells were used in downstream analyses.

### Statistical analysis

#### Calculation of size factor normalized cell class counts

Size factor normalized cell counts were calculated as in^28, 29^. Briefly, counts for cells of the same cell class within each organoid were added up and organized into a cell class by organoid matrix. A Monocle3 cds was created from the matrix, and *`size factors()`*, which takes the total number of cells in the organoid and divides it by the geometric mean of cell counts for all organoids, was run. The number of cells of each cell class in each organoid was divided by the size factor of that organoid. All values were rounded to the nearest whole number.

#### Detection of significant fold changes in cell class abundance

A generalized linear model was fit to the normalized data using a beta-binomial distribution^65^ as in^28, 29^.

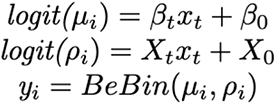

Where *y_i_* represents the normalized cell count distribution for cell class *i*, *μ_i_* is the mean of the distribution, and ϱ*_i_* is the overdispersion or “litter effect” associated with the beta-binomial distribution. The binary indicator variable *X_t_* denotes which treatment the organoids were exposed to. All D28 treatments were first modeled with respect to the None treatment. The CHIR, SU5402, and SU5402:CHIR treatments were later modeled compared to the BMP treatment. At D63 CHIR and SU5402:CHIR treatments were modeled compared to BMP. The models were fit using the VGAM package, and p-values were determined by a Wald test on β_t_^66, 67^. Abundance changes were determined significant if the q-value after Benjamini Hochberg correction was < 0.05^68^. In D28 organoids, the model above was also used to determine significant abundances changes in specific retinal cell populations. When this was performed, all cell classes were included in the calculation even though only the retinal cell classes are displayed in the heatmap in Figure S3D.

#### Clustering of individual organoids

Counts for cells of the same cell class within each organoid were added up and organized into a cell class by organoid matrix. A Monocle3 cds was created from the matrix and `size factors()` was run. The cds was processed `preprocess_cds(cds, method = “PCA”, norm_method = “size_only”, num_dim = 9)` and then subjected to dimensionality reduction in two dimensions by UMAP `reduce_dimension(cds, umap.n_neighbors = 30, umap.min_dist = 0.05)`. Clustering was performed with a resolution of 1x10^-2^.

#### Detection of treatments and cell clusters with high variance

A cluster by individual organoid matrix was calculated, size factors were determined, and a generalized linear model was fit to the normalized data using a beta-binomial distribution as above. The VGAM package doesn’t automatically model dispersions for the beta-binomial distribution, so `simulate.vlm()` was run 100 times to model the mean and standard deviation which was then used to calculate the coefficient of variation (*CV = σ/μ*) for each cluster and each treatment. A gamma-valued glm of the form pioneered by DESeq^69^ was then to capture the trend between the average number of cells in a cluster across organoids and that cluster’s CV using VGAM^66, 67^. To identify clusters with higher than expected CV, conditional upon treatment, we tested whether the CV for a given cluster exceeded this model’s 95% confidence interval’s upper limit. To assess whether a treatment altered heterogeneity compared to the BMP treatment, a likelihood-ratio test was used to compare the output of the gamma distribution to a reduced model that excludes the treatment term. Treatments were considered more heterogeneous than BMP if the likelihood-ratio test produced a q-value after Benjamini Hochberg correction of < 0.05 indicating that use of `treatment` as a factor improves the model and the sign of the treatment term was positive indicating that the treatment-dependent change led to an increase in heterogeneity^68^.

## Supplemental Information

**Table S1. Cost benefit analysis of sci-Plex**

Demonstration of the cost savings of sci-Plex over 10x sequencing

**Table S2. Cell class marker genes**

A list of the genes used to assign cell classes for all sci-Plex experiments

**Table S3. Variance modeling output**

The coefficients and likelihood ratio test results obtained after modeling the CV vs cluster size with a gamma distribution. The likelihood test compared CHIR or SU5402:CHIR to BMP treatments.

## Data and code availability

All single cell datasets have been uploaded and are publicly available at NCBI’s GEO database with accession number GSE220661. The code used to process the output from sci-Plex experiments is available at the Brotman Baty Institute’s github page (https://github.com/bbi-lab) under the bbi-sci and bbi-dmux repositories. Custom code will be made publicly available.

