## Supplemental Figures for "Single-cell sequencing of individual retinal organoids reveals determinants of cell fate heterogeneity"

### Supplemental Information

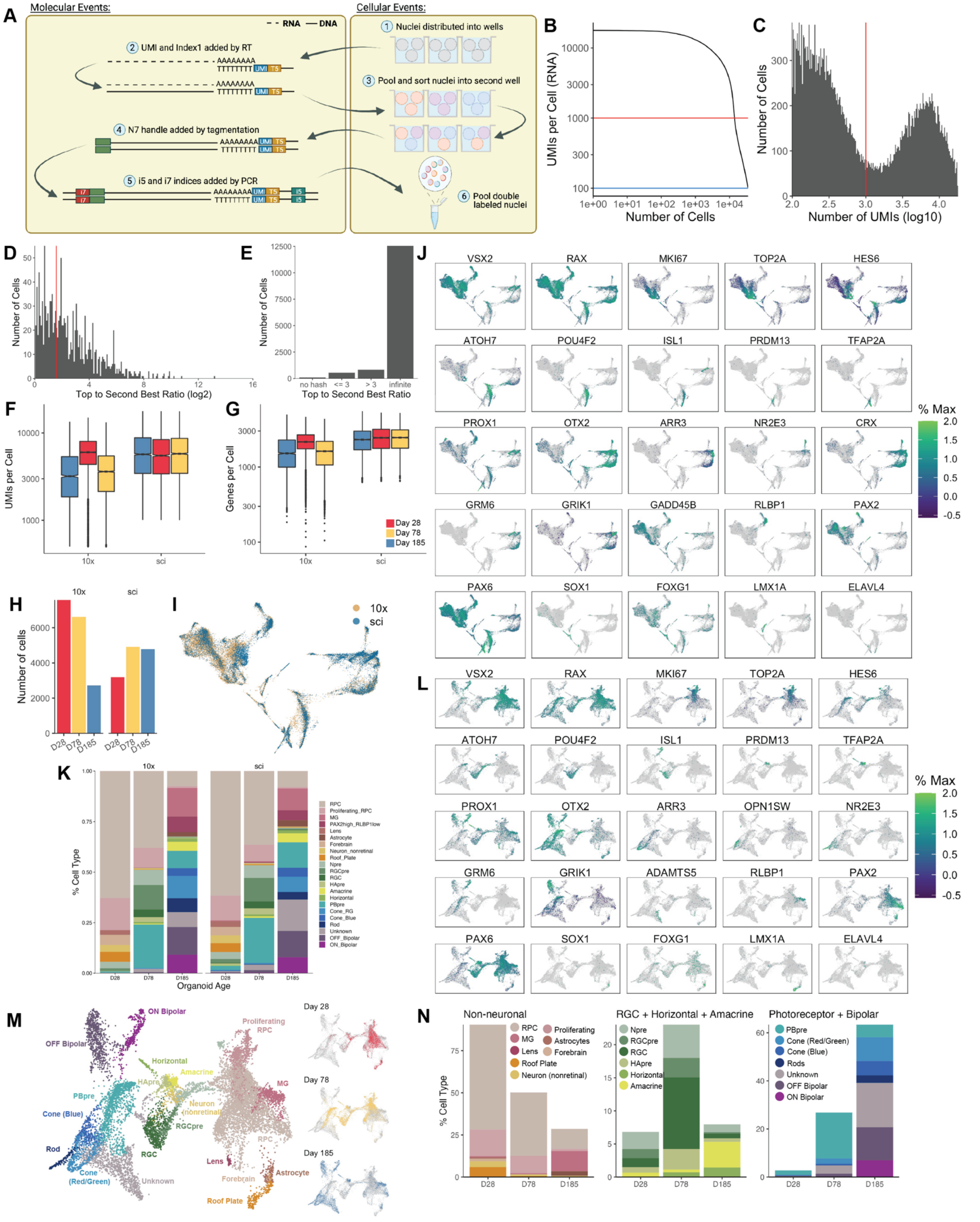
A
Molecular Events:
Cellular Events:
RNA
DNA
1 Nuclei distributed into wells
2 UMI and Index1 added by RT
3 Pool and sort nuclei into second well
4 N7 handle added by tagmentation
5 i5 and i7 indices added by PCR
6 Pool double labeled nuclei
B
UMIs per Cell (RNA)
Number of Cells
C
Number of Cells
Number of UMIs (log10)
D
Number of Cells
Top to Second Best Ratio (log2)
E
Number of Cells
no hash
<= 3
> 3
infinite
Top to Second Best Ratio
F
UMIs per Cell
10x
sci
G
Genes per Cell
Day 28
Day 78
Day 185
H
Number of cells
I
10x
sci
J
VSX2
RAX
MKI67
TOP2A
HES6
ATOH7
POU4F2
ISL1
PRDM13
TFAP2A
PROX1
OTX2
ARR3
NR2E3
CRX
GRM6
GRIK1
GADD45B
RLBP1
PAX2
PAX6
SOX1
FOXG1
LMX1A
ELAVL4
% Max
K
% Cell Type
Organoid Age
L
OPN1SW
ADAMTS5
M
ON Bipolar
OFF Bipolar
Horizontal
Amacrine
HApre
PBpre
Cone (Blue)
Rod
Cone (Red/Green)
Unknown
RGC
RGCpre
Neuron (nonretinal)
Proliferating RPC
MG
RPC
Lens
Forebrain
Astrocyte
Roof Plate
Day 28
Day 78
Day 185
N
Non-neuronal
RGC + Horizontal + Amacrine
Photoreceptor + Bipolar
% Cell Type
Proliferating
Astrocytes
Neuron (nonretinal)
Npre
Cone (Red/Green)
Rods
OFF Bipolar

**Figure S1. Quality control of sci-Plex and 10x comparison datasets. A)** sci-RNA-seq relies on successive split-pool barcoding of RNA: permeabilized nuclei are distributed into wells of a 96-well plate. Reverse transcription is performed using a polyD-T oligo that contains both a UMI and barcode (T5). Each well is provided a unique oligo such that all cDNA molecules produced in a well will have the same barcode. Cells are then pooled and redistributed into a second set of 96-well plates. The cDNA is converted to dsDNA and subjected to tagmentation which adds a uniform sequence to the end of the cDNA molecules. This sequence and the sequence at the end of the RT oligos can be used to amplify the library by PCR. The PCR primers each contain barcode sequences (i7, i5) and the wells all receive a unique combination of PCR oligos. Experiments can be designed such that the vast majority of cells travel through a unique set of wells. Thus, RNAs that share the same barcode combination are assumed to have come from the same cell. **B)** A knee plot of the number of UMIs per cell compared to the number of cells recovered. The red line (1000UMIs) indicates the value denoting whether a barcode combination should be considered a cell. **C)** A histogram of the number of cells per UMI count. UMI counts left of the red line (1000 UMIs) are removed from the analysis. **D)** Hashes were sequenced and assigned to cells. A Top to Second Best Ratio was then calculated by dividing the UMI count of the most common hash barcode by the UMI count of the second most common hash barcode detected for an individual cell. A histogram of the number of cells recovered at each Top to Second Best Ratio was then generated. The red line (3) denotes the Top to Second Best Ratio required for a cell to be assigned a confident hash call. **E)** A bar plot of the number of cells with Top to Second Best Ratios matching given criteria. In the event only a single hash barcode is detected, the ratio is infinite. **F)** Bar plots of the number of cells recovered for each sample. **G)** Box plots of the number of UMIs recovered from each cell for each sample. **H)** Box plots of the number of genes recovered from each cell for each sample. **I)** Integration of 10x and sci data by Seurat's MNN-CCA methodology. The cells are colored by the technology with which they were prepared for sequencing. **J)** Feature plots of the genes used to define cell types from the integrated sci-Plex and 10x UMAP. **K)** Stacked bar plots demonstrating the cell type compositions of organoids from D28, D78, and D185 organoids as prepared for sequencing by 10x and sci-Plex. RPC: retinal progenitor cell, MG: Müller glia, Npre: retinal neuronal precursor, RGC: retinal ganglion cell, RGCpre: RGC precursor, HApre: horizontal/amacrine precursor, PBpre: photoreceptor/bipolar precursor, Cone_RG: red/green cone. **L)** A UMAP was generated by Monocle3 using only the cells recovered by sci-Plex. Cells are colored by expression of the indicated gene. The displayed genes were used when defining cell types. **M)** The UMAP from L). Left: Cells are colored by cell type, Right: Cells are faceted and colored by organoid age. **N)** Using the cell type assignments from M, cell types were split into Brain/Non-neuronal (Non-neuronal), RGC+Horizontal+Amacrine (RGC+H+A), and Photoreceptor+Bipolar (PR+BP) categories. Cells were counted and the % Cell Type was determined for each organoid age.

—-------------------------------------

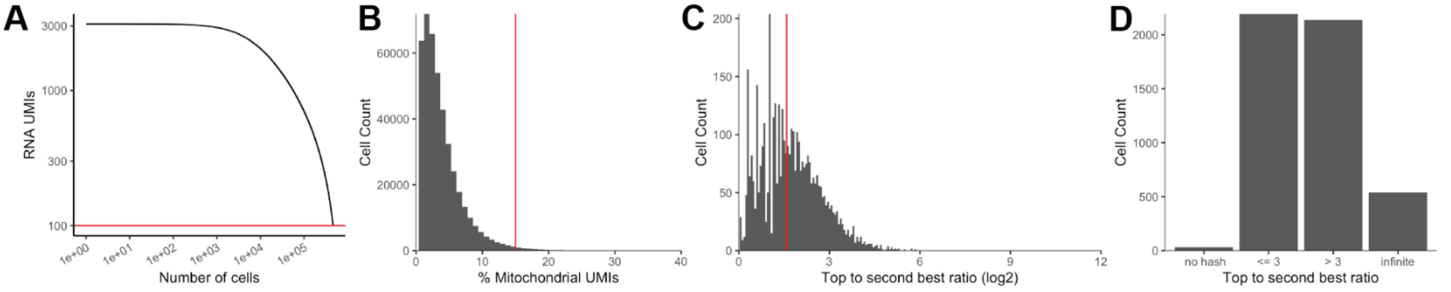

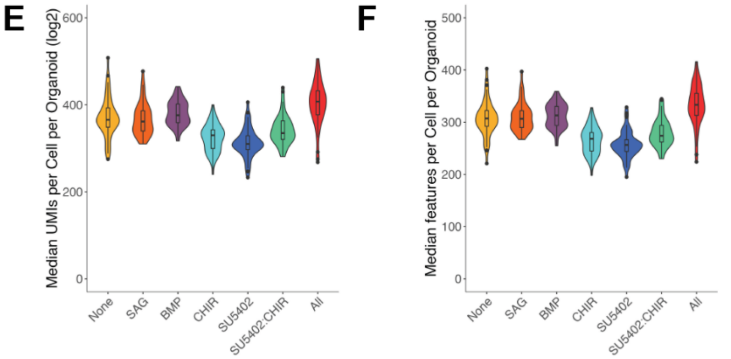

G

| sci-Plex oligo plate | Condition 1 | Condition 2 |
|---|---|---|
| 1 | None | SAG |
| 2 | SU5402 | CHIR |
| 3 | SU5402:CHIR | BMP |
| 4 | All | |

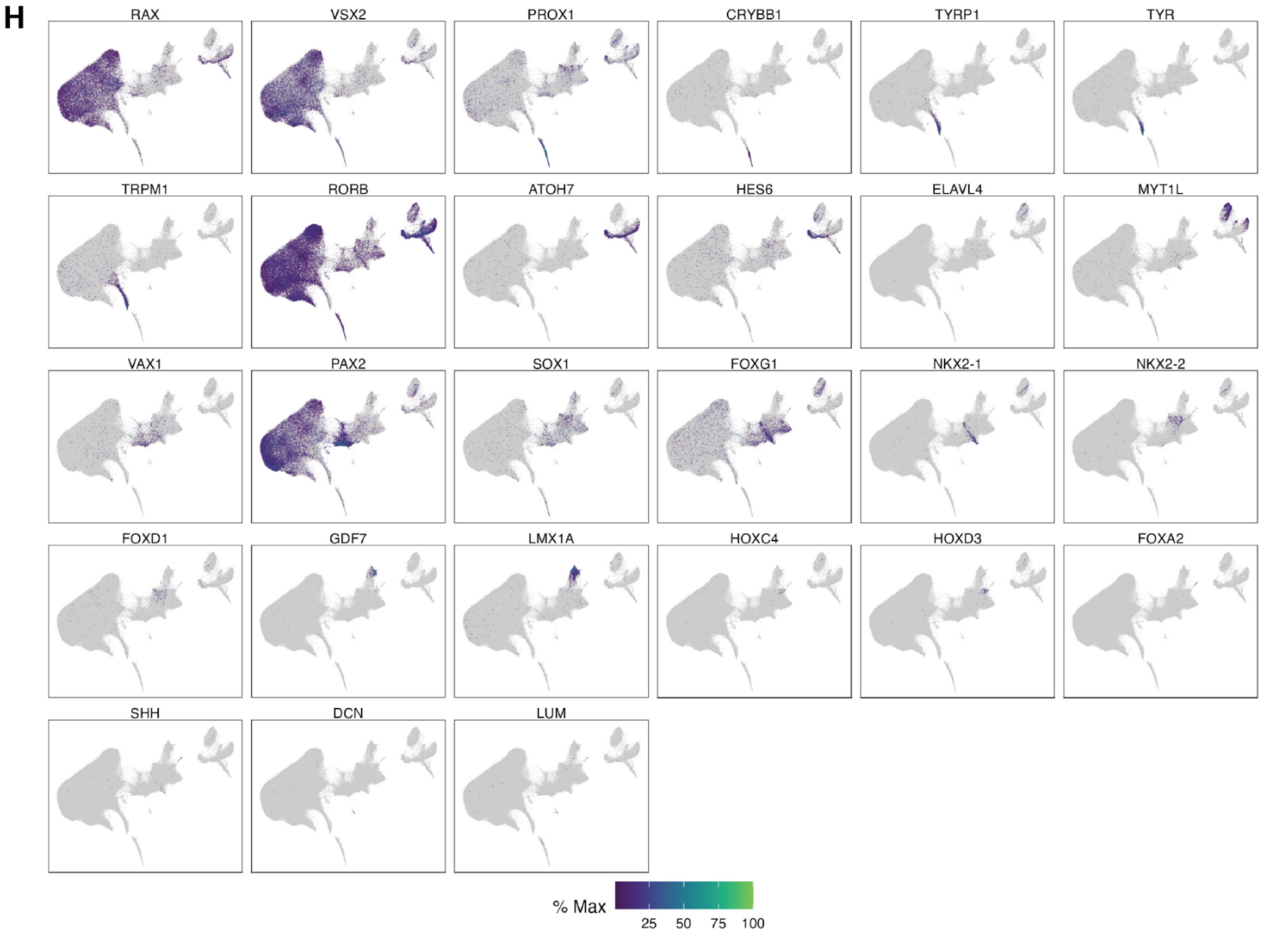

**Figure S2. Quality control of sci-Plex performed on individual D28 organoids. A)** A knee plot of the number of UMIs per cell compared to the number of cells recovered from D28 organoids. The red line (100 UMIs) indicates the value denoting whether a barcode combination should be considered a cell. **B)** A histogram of the % mitochondrial UMIs. Cells with percentages right of the red line (15%) are removed from the analysis. **C)** A histogram of the number of cells recovered at each Top to Second Best Ratio was then generated. The red line (2.5) denotes the Top to Second Best Ratio required for a cell to be assigned a confident hash call. In the event only a single hash barcode is detected, the ratio is infinite. **D)** A bar plot of the number of cells with Top to Second Best Ratios matching given criteria. **E)** Violin plots of the number of UMIs recovered from each cell for each D28 organoid per treatment. **F)** Violin plots of the number of features recovered from each cell for each D28 organoid per treatment. **G)** Table indicating which sci-Plex oligo plate was used to hash each of the treatment conditions. **H)**Expression plots of the genes used to define the cell types in Figure 2E.

—-------------------------------------

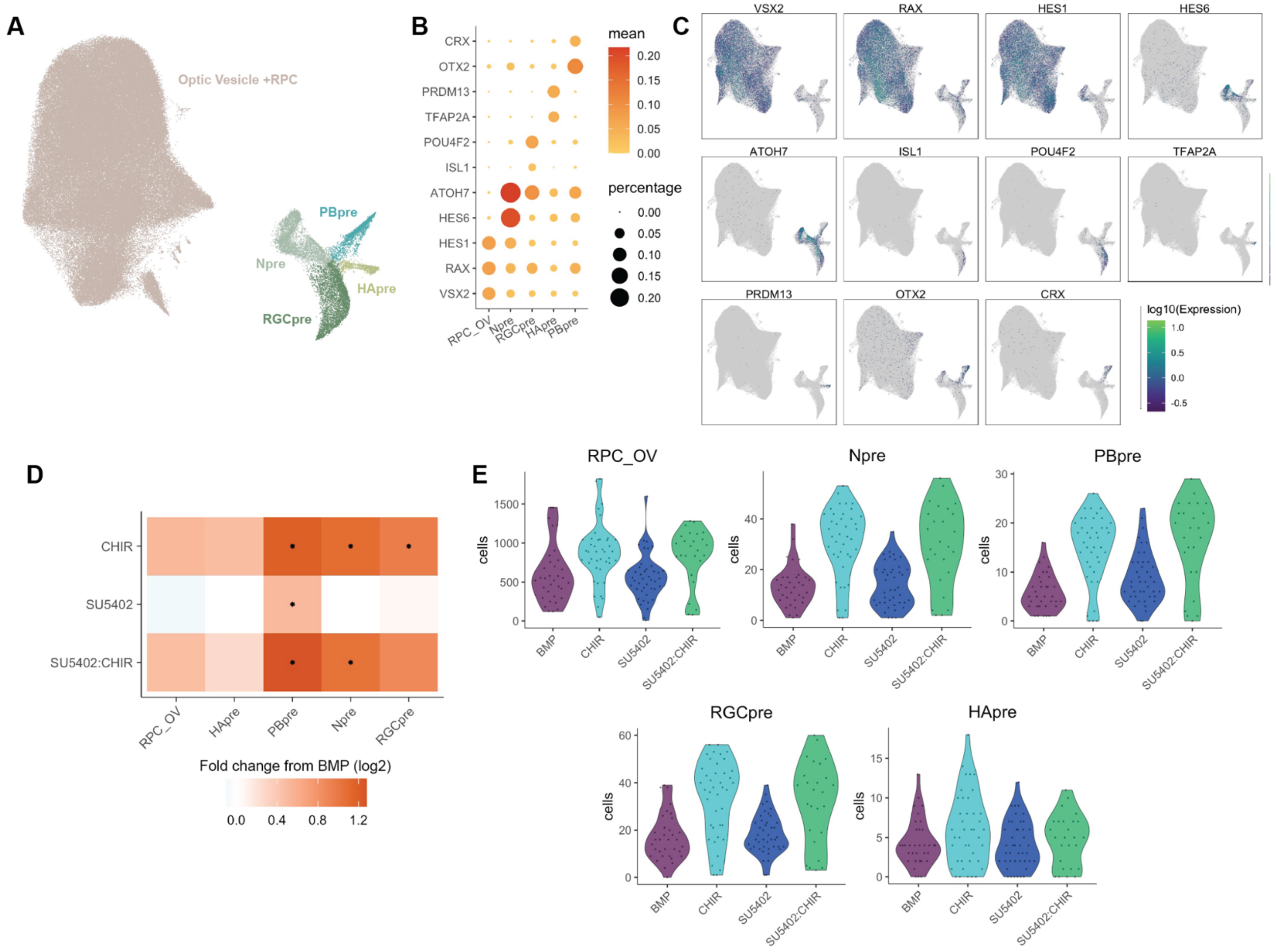

**Figure S3. Investigation of abundance changes in specific retinal cell types. A)** A UMAP of the Retina and RPC_OV subset of cells from D28 organoids was derived by Monocle3. Cells are colored by retinal cell type. RPC_OV: retinal progenitor cell/optic vesicle, Npre: retinal neuronal precursor, RGCpre: retinal ganglion cell precursor, HApre: horizontal/amacrine precursor, PBpre: photoreceptor/bipolar precursor. **B)** A dot plot of the genes used to identify the retinal cell types in A. **C)** Expression plots of the genes used to define the cell types in Figure S4A. **D)** Heatmap of fold change in cell type abundance compared to “BMP” for D28 organoids. A beta-binomial model was fit to the data and used to determine whether there were significant changes in cell type abundances. A * in the center of the box indicates a q-value of less than 0.05 after Benjamini Hochberg correction. Note that while only the retinal cell types are displayed, the test was performed on size-factor normalized cell counts of all cell types in the organoid. **E)** Violin plots of size-factor normalized cell counts for the retinal cell types. Plots are colored by treatment.

—-----------------------------------------

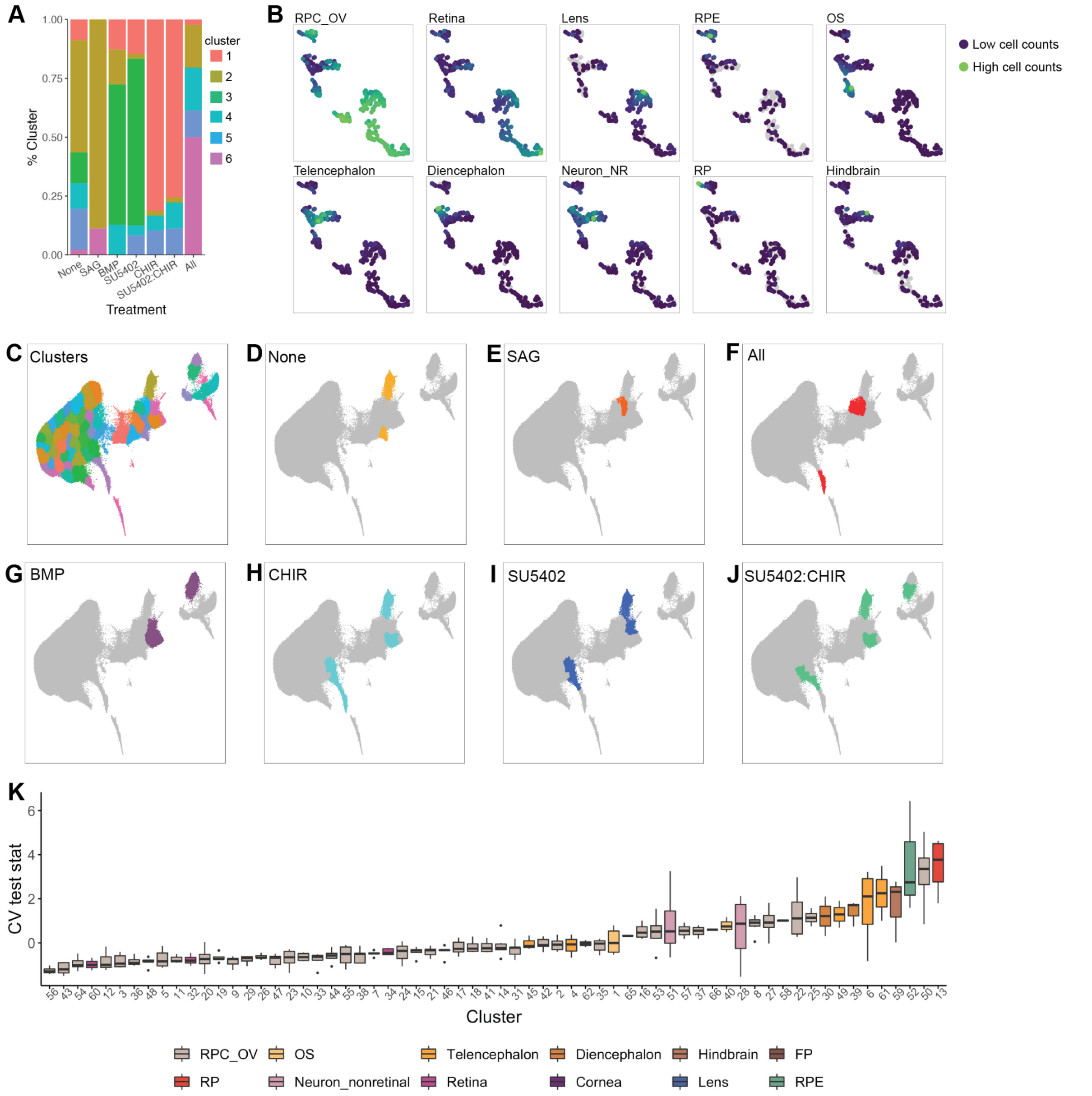

**Figure S4. Identification of cell types with high variability in D28 organoids. A)** Stacked bar plot demonstrating the distribution of organoids across archetypes. **B)** UMAP from 4J in which color indicates the abundance of each cell type. Each dot represents an individual organoid. Blue indicates few cells of that type are found in that organoid and green indicates high counts for that cell type. RPC_OV: retinal progenitor cell/optic vesicle, RPE: retinal pigmented epithelium, OS: optic stalk, Neuron_NR: non-retinal neuron, RP: roof plate. **C)** UMAP from 2C of the individual clusters used for analyses modeling the variance across treatments and clusters. **D)** UMAP highlighting the clusters with significantly higher variance than expected in None, **E)**

SAG, **F)** All, **G)** BMP, **H)** CHIR, **I)** SU5402, and **J)** SU5402:CHIR. **K)** Box plots with clusters ordered according to increasing CV test stat. A single CV test stat is calculated for each treatment, and the boxplots summarize the CV test stat values across treatments for each cluster. Box plots are colored by the cell type assignment as in Figure 2E. RPC_OV: retinal progenitor cell/optic vesicle, RPE: retinal pigmented epithelium, OS: optic stalk, RP: roof plate, FP: floor plate.

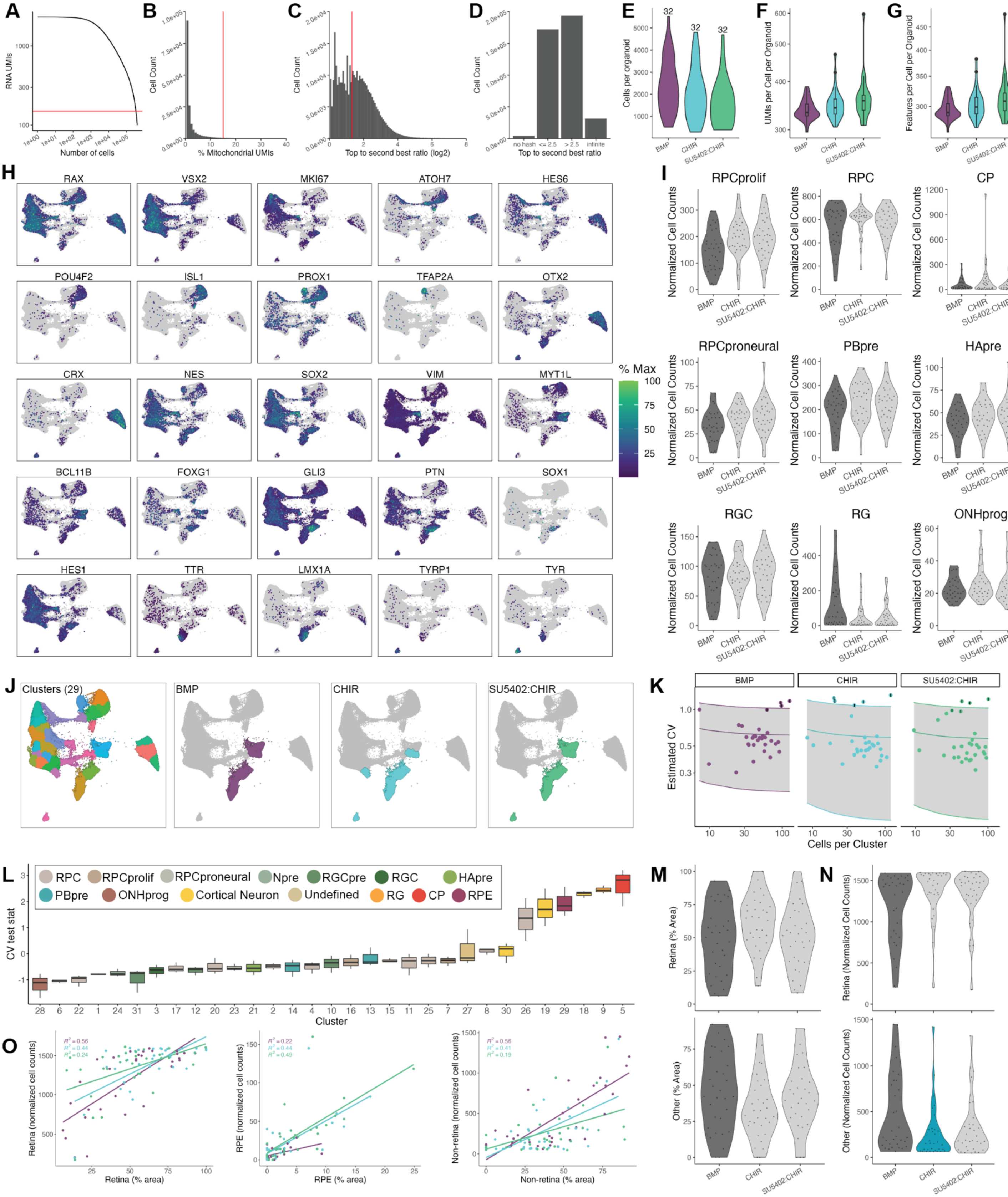
A
RNA UMIs
Number of cells
B
Cell Count
% Mitochondrial UMIs
C
Cell Count
Top to second best ratio (log2)
D
Cell Count
no hash
<= 2.5
> 2.5
infinite
Top to second best ratio
E
Cells per organoid
32
32
32
F
UMIs per Cell per Organoid
G
Features per Cell per Organoid
BMP
CHIR
SU5402:CHIR
H
RAX
VSX2
MKI67
ATOH7
HES6
POU4F2
ISL1
PROX1
TFAP2A
OTX2
CRX
NES
SOX2
VIM
MYT1L
BCL11B
FOXG1
GLI3
PTN
SOX1
HES1
TTR
LMX1A
TYRP1
TYR
% Max
I
RPCprolif
RPC
CP
RPCproneural
PBpre
HApre
RGC
RG
ONHprog
Normalized Cell Counts
J
Clusters (29)
BMP
CHIR
SU5402:CHIR
K
Estimated CV
Cells per Cluster
L
RPC
RPCprolif
RPCproneural
Npre
RGCpre
RGC
HApre
PBpre
ONHprog
Cortical Neuron
Undefined
RG
CP
RPE
CV test stat
Cluster
M
Retina (% Area)
Other (% Area)
N
Retina (Normalized Cell Counts)
Other (Normalized Cell Counts)
O
Retina (normalized cell counts)
Retina (% area)
RPE (normalized cell counts)
RPE (% area)
Non-retina (normalized cell counts)
Non-retina (% area)

**Figure S5. Quality control of the D63 organoid datasets A)** A knee plot of the number of UMIs per cell compared to the number of cells recovered. The red line (150 UMIs) indicates the value denoting whether a barcode combination should be considered a cell. **B)** A histogram of the number of cells per % mitochondrial UMIs. All cells with > 15% were discarded. **C)** A histogram of the number of cells recovered at each Top to Second Best Ratio was then generated. The red line (2.5) denotes the Top to Second Best Ratio required for a cell to be assigned a confident hash call. **D)** A bar plot of the number of cells with Top to Second Best Ratios matching given criteria. In the event only a single hash barcode is detected, the ratio is infinite. **E)** Violin plot of the number of cells recovered for each organoid across treatments. **F)** Violin plot of the number of UMIs recovered from each cell per D63 organoid per treatment. **G)** Violin plot of the number of features recovered from each cell per D63 organoid per treatment. **H)** Expression plots of the genes used to define cell types in Figure 5A. **I)** Violin plots of size-factor normalized cell counts for D63 organoids. The dark gray indicates the treatment that was used for fold-change calculations in 5B, and light gray indicates no significant change in cell type abundance. **J)** A beta-binomial distribution was used to determine the dispersions across treatments for each of the 29 clusters used to define cell types. The mean and CV calculated from the beta-binomial distribution, were then used to model the relationship between the number of cells per cluster and the CV using a gamma distribution. Left: UMAP with the cells colored by the 29 clusters used to define cell types. Right: UMAPs in which the clusters highlighted are more variable across treatments than expected by chance. Cells are colored by treatment. **K)** Displayed are the modeled relationships between mean cell count and CV per cluster colored by treatment. **L)** Boxplots with clusters ordered according to increasing CV test stat. A single CV test stat is calculated for each treatment, and the boxplots summarize the CV test stat values across treatments. RPC: retinal progenitor cell, RPCprolif: proliferating RPC, RPCproneural: proneural RPC, Npre: retinal neural precursor, RGC: retinal ganglion cell, RGCpre: RGC precursor, HApre: horizontal/amacrine precursor, PBpre: photoreceptor/bipolar precursor, ONHprog: optic nerve head progenitor, RG: radial glia, CP: choroid plexus, RPE: retinal pigmented epithelium. **M)** Violin plots of % Area measurements for Retina and Non-retinal (Other) cell types as calculated from bright phase microscopy images of D62 organoids. Plots are colored as in S6H. **N)** Violin plots of size-factor normalized Retina and Non-retinal (Other) cell counts for D63 organoids using broad cell types. Plots are colored as in S6H with teal indicating a significantly decreased cell type abundance. **O)** Scatter plots showing the correlation between the sci-Plex cell counts and % Area for each individual D63 organoid across treatments. $R^2$ values are in the top right corner colored by treatment.

—------------------------------------

**Table S1. Cost benefit analysis between sci-Plex and 10x***

| Experiment | Total cells | sci-Plex cost per cell | 10x cost per cell | sci-Plex cost per sample prep | 10x cost per sample prep |
|---|---|---|---|---|---|
| 10x match | 16,914 (10x) 12,867 (sci) | $0.14 | $0.35** | $608 | $2000* |
| D28 | 203,511 | $0.04 | $0.30*** | $23 | $196*** |
| D63 | 194,335 | $0.11 | $0.0625*** | $76 | $232*** |

* This analysis does not take into account the sequencing cost. However, sci usually results in fewer UMIs which can make it difficult for some gene expression analyses, but it does make it easier to sequence for a lower cost.

**This was the actual cost for the experiment as performed. Each age was loaded onto its own lane with a desired recovery of 7000 cells per well.

***This estimate assumes multiplexing of 16 samples per lane using the CellPlex kit. A desired recovery of 500 cells per sample (8000 cells per lane) was assumed for the D28 experiment and 2000 cells per sample for the D28 experiment (32,000 cells per lane). These values are based off of the median number of cells recovered per organoid from the sci-Plex experiments.

**Table S2. Cell type marker genes**

A list of the genes used to assign cell types for all sci-Plex experiments

| Cell Type | Expt | Enriched Markers | | | |
|---|---|---|---|---|---|
| RPC | 10x_sci comp | VSX2 | RAX | | |
| Proliferating RPC | 10x_sci comp | VSX2 | RAX | MKI67 | TOP2A |
| MG | 10x_sci comp | RLBP1 | | | |
| PAX2high RLBP1low | 10x_sci comp | PAX2 | RLBP1 | | |
| Lens | 10x_sci comp | PROX1 | CRYBB1 | | |
| Astrocyte | 10x_sci comp | PAX2 | GFAP | | |
| Forebrain | 10x_sci comp | SOX1 | FOXG1 | | |
| Neuron nonretinal | 10x_sci comp | ELAVL4 | DLX6 | MYT1L | |
| Roof plate | 10x_sci comp | GDF7 | LMX1A | | |
| Npre | 10x_sci comp | HES6 | ATOH7 | | |
| RGCpre | 10x_sci comp | ATOH7 | POU4F2 | | |

| | | | | | | | |
|---|---|---|---|---|---|---|---|
| RGC | 10x_sci comp | POU4F2 | ISL1 | | | | |
| HApre | 10x_sci comp | PRDM13 | TFAP2A | | | | |
| Amacrine | 10x_sci comp | TFAP2A | | | | | |
| Horizontal | 10x_sci comp | PROX1 | TFAP2A | ONECUT1 | | | |
| PBpre | 10x_sci comp | HES6 | OTX2 | CRX | | | |
| Cone RG | 10x_sci comp | ARR3 | CRX | OTX2 | | | |
| Cone Blue | 10x_sci comp | OPN1SW | CRX | OTX2 | | | |
| Rod | 10x_sci comp | NRL | CRX | OTX2 | | | |
| Unknown | 10x_sci comp | PIK3CG | PP1R27 | GDF15 | KBTBD12 | INPP5D | OTX2 |
| OFF Bipolar | 10x_sci comp | VSX1 | GRIK1 | OTX2 | | | |
| ON Bipolar | 10x_sci comp | VSX2 | ISL1 | PROX1 | GRM6 | OTX2 | |
| RPC/OV | D28 All cells | RAX | VSX2 | HES1 | | | |
| OS | D28 All cells | VAX1 | PAX2 | SOX1 | | | |
| Telencephalon | D28 All cells | SOX1 | FOXG1 | NKX2-1 | | | |
| Diencephalon | D28 All cells | SOX1 | FOXD1 | NKX2-2 | | | |
| Hindbrain | D28 All cells | HOXC4 | HOXD3 | HOXB6 | | | |
| FP | D28 All cells | FOXA2 | SHH | | | | |
| RP | D28 All cells | GDF7 | TTR | LMX1A | | | |
| Neuron nonretinal | D28 All cells | ELAVL4 | MYT1L | EOMES | FOXG1 | | |
| Retina | D28 All cells | RORB | ATOH7 | ONECUT2 | HES6 | RAX | |
| Cornea | D28 All cells | DCN | LUM | | | | |
| Lens | D28 All cells | PROX1 | CRYBB1 | | | | |
| RPE | D28 All cells | TYRP1 | TYR | TRPM1 | OTX2 | | |
| RPC/OV | D28 Retina | RAX | VSX2 | HES1 | | | |
| Npre | D28 Retina | HES6 | ATOH7 | | | | |
| RGCpre | D28 Retina | ATOH7 | POU4F2 | ISL1 | | | |
| HApre | D28 Retina | TFAP2A | PRDM13 | | | | |
| PBpre | D28 Retina | OTX2 | CRX | | | | |
| RPC | D63 | RAX | VSX2 | RORB | | | |

| | | | | | | | | | | |
|---|---|---|---|---|---|---|---|---|---|---|
| RPCprolif | D63 | RAX | VSX2 | MKI67 | TOP2A | RORB | | | | |
| RPCproneural | D63 | MKI67 | TOP2A | ATOH7 | HES6 | RORB | ONECUT1 | | | |
| Npre | D63 | ATOH7 | HES6 | RORB | ONECUT1 | | | | | |
| RGCpre | D63 | ATHO7 | POU4F2 | RORB | ONECUT1 | | | | | |
| RGC | D63 | POU4F2 | ISL1 | RORB | ONECUT1 | | | | | |
| HApre | D63 | PROX1 | TFAP2A | PRDM13 | RORB | ONECUT1 | | | | |
| PBpre | D63 | OTX2 | CRX | ONECUT1 | RORB | | | | | |
| ONHprog | D63 | NES | SOX2 | VIM | SOX1 | RAX | VSX2 | MKI67 | TOP2A | RORB |
| Transition | D63 | N/A | | | | | | | | |
| Cortical Neuron | D63 | MYT1L | BCL11B | | | | | | | |
| Radial Glia | D63 | GLI3 | FOXG1 | SOX1 | PTN | SOX2 | HES1 | PPRX1 | OTX1 | |
| Choroid Plexus | D63 | TTR | LMX1A | WNT2B | WNT3A | OTX2 | | | | |
| RPE | D63 | TYRP1 | TYR | OTX2 | MITF | | | | | |

**Table S3. Variance modeling output**

The coefficients and likelihood ratio test results obtained after modeling the CV vs cluster size with a gamma distribution. The likelihood test compared CHIR or SU5402:CHIR to BMP treatments.
